# Identification of *SNARE* Genes in Cucumber and the Role of *CsSYP121* in Salt Stress Response

**DOI:** 10.64898/2026.03.30.715073

**Authors:** Weijuan Zhou, Jie Zheng, Shengjun Zhou, Yue Guo, Dongmei Kong, Pu Yang, Ben Zhang

## Abstract

Soluble N-ethylmaleimide-sensitive factor attachment protein receptors (SNAREs) are essential regulators of plant growth, development, and stress adaptation. In this study, we performed a comprehensive genome-wide identification of *SNARE* genes in cucumber (*Cucumis sativus* L.), uncovering 51 putative members designated as *CsSNAREs*. Phylogenetic analysis confirmed that these genes cluster into five major clades: *Qa-CsSNARE* (14), *Qb-CsSNARE* (9), *Qc-CsSNARE* (10), *Qb+c-CsSNARE* (3), and *R-CsSNARE* (15). Bioinformatic analysis of their promoter regions, coupled with expression profiling under diverse abiotic stress conditions, highlighted a heightened responsiveness within the *Qa-CsSNARE* subfamily. To validate this, we selected representative *Qa-CsSNARE* genes for quantitative real-time PCR analysis under drought and salt stress. Among these, *CsSYP121* was notably induced by salt treatment. We subsequently generated transgenic cucumber lines overexpressing *CsSYP121* and challenged them with salinity. Phenotypic assessment, combined with measurements of reactive oxygen species (ROS) accumulation and K^+^/Na^+^ ratios, demonstrated that *CsSYP121* overexpression (OE) confers enhanced salt tolerance and boosts antioxidant capacity. We propose a model wherein *CsSYP121* mitigates ROS-induced cellular damage under salt stress, potentially through promoting K^+^/Na^+^ homeostasis, thereby improving plant performance under saline conditions. Our findings identify *CsSYP121* as a promising candidate gene for breeding salt-tolerant crops.

## 1. Introdution

Interorganellar communication in plant cells relies critically on membrane integrity and functional dynamics. This communication regulates cellular homeostasis, coordinated growth, and developmental transitions (Gu et al., 2020; Kwon et al., 2020; Zhang et al., 2020). The soluble N-ethylmaleimide-sensitive factor attachment protein receptor (SNARE) protein family are known to be involved in vesicle trafficking through driving membrane fusion (Hong, 2005). SNARE proteins typically range in size from 100 to 300 amino acids and often contain at least one region approximately 60 amino acid residues long, called SNARE motif. These proteins interact with each other to form SNARE complexes, which mediate vesicular fusion. Based on their subcellular localization, SNARE proteins can be classified into t (target)-SNAREs and v (vesicle)-SNAREs. In addition, SNARE members exhibit distinct classification based on amino acid residues within their central layer, leading to the designation of Q (glutamine-rich) and R (arginine-rich) sub-families (Fasshauer et al., 1998). The Q subgroup includes Qa-, Qb-, Qc-, and Qb+c-SNAREs. Structurally, most Q-SNAREs have an N-terminal autoinhibitory Habc domain (triple helical motifs Ha, Hb, Hc), a core SNARE motif, and a C-terminal transmembrane domain (Zhang et al., 2020). Vesicle fusion typically requires three Q-SNAREs and one R-SNARE (Cermesoni et al., 2024; Zhang et al., 2020).

SNARE proteins function in key plant processes, including phytohormone signaling (Xia et al., 2020; Xue et al., 2018), pathogen defense (Waghmare et al. 2018; Zhu et al. 2024), stress adaptation (Kwon et al., 2020; Nisa et al., 2017; Waghmare et al., 2024), gravity perception (Yano et al., 2003), cell plate assembly (Zhang et al., 2011), and arbuscular mycelium morphogenesis (Huisman et al., 2016; Sogawa et al., 2019).

SNARE genes are identified in many plants, including Arabidopsis, wheat (*Triticum aestivum*), rice (*Oryza sativa*), Chinese cabbage (*Brassica rapa*), grapevine (*Vitis vinifera*), *Brassica napus*, and foxtail millet (*Setaria italica*) (Bassham et al., 2008; Zhang et al., 2018; G. Wang et al., 2021; Xu et al., 2022; H. Wang et al., 2021; Zeng et al., 2024). Plants possess more *SNARE* genes than animals or microorganisms, suggesting roles in complex plant physiology. In *Marchantia polymorpha*, some SNARE localizations differ from Arabidopsis homologs; for example, Qa-SNAREs MpSYP12A and MpSYP12B localize to oil bodies involved in lipid metabolism (Kanazawa et al., 2016). Some Qa- and R-SNAREs also regulate membrane transporters or ion channels via direct interactions, affecting ion homeostasis. This is crucial for plant growth and abiotic stress responses, especially to drought and salinity (Grefen et al., 2015; Xia et al., 2019; Zhang et al., 2022, 2020, 2015).

Qa-SNARE SYP121 and SYP122 are conserved in vegetative plants (Sanderfoot 2007; Waghmare et al. 2018). They share high sequence similarity and form a SNARE complex with Qb+c-SNARE SNAP33 and R-SNAREs VAMP721. This complex mediates vesicle trafficking to the plasma membrane, linked to biotic stress responses (Karnik et al. 2013; Waghmare et al. 2018; Enami et al. 2009; Kwon et al. 2020, 2008). In Arabidopsis, single *syp121* or *syp122* mutations show minimal effects, but double mutations are lethal, indicating functional redundancy (Karnik et al., 2017). However, SYP121 and SYP122 mediate distinct secretory pathways (Waghmare et al. 2018). Recent research reveals that SYP121 physically interacts with potassium channels KAT1 and KC1, regulating their gating and linking vesicle trafficking to potassium homeostasis (Grefen et al., 2010a, 2015). SEC11, a Sec1/Munc18 protein, selectively interacts with SYP121 via R^20^R^21^ residues to regulate SNARE-channel complex formation (Zhang et al., 2019). These findings highlight SYP121’s critical role in plant growth and development.

Cucumber, *Cucumis sativus* L., is a nutrient-dense, low-calorie vegetable rich in vitamin C, fiber, and water, is globally significant (Huang et al., 2009). Cucumber is one of the most water-loving vegetable crop, it is sensitive to salt and drought stress (Das et al., 2024; Li et al., 2023; Wang et al., 2018). However, studies on cucumber *SNARE* genes are limited. The recent near-complete cucumber genome (Guan et al., 2024), enabling systematic investigations into cucumber genes, particularly the *CsSNARE* genes.

In this research, we conducted a systematic screening and successful identification of the *CsSNARE* gene family within the cucumber genome. Furthermore, the expression profiles of key *Qa-CsSNARE* genes were rigorously characterized following exposure to both drought and salt stresses, thereby establishing a foundational understanding necessary for elucidating their potential contributions to abiotic stress tolerance mechanisms. To probe the role of individual members, we developed *CsSYP121*-overexpressing cucumber lines and evaluated their performance, including growth, development, and physiological responses under salt stress. These results offer valuable molecular insights for future functional studies aimed at enhancing crop resilience and provide new genetic resources for improving salt tolerance in cucumber breeding programs.

## 2. Materials and methods

### 2.1 Identification of the CsSNARE gene family

The Cucumber (Chinese Long) v4 genome dataset was obtained from the Cucumber Genome Database (http://www.cucumberdb.com, accessed on 05 May 2025) (Guan et al., 2024). Candidate *CsSNARE* genes identified through BLAST analysis of Arabidopsis SNARE sequences (retrieved from The Arabidopsis Information Resource, https://www.arabidopsis.org, accessed on 05 May 2025) in the cucumber genome. The truncated open reading frames, peptides <100 amino acids, and redundant sequences were removed. For the validated CsSNARE proteins, theoretical isoelectric points (pI), molecular weights (MW), and amino acid compositions were predicted using ExPASY (http://web.expasy.org/protparam, accessed on 03 June, 2025) (Gasteiger, 2003) .

### 2.2 Phylogenetic analysis and chromosomal localization of the cucumber CsSNARE gene family

For phylogenetic analysis, full-length SNARE protein sequences from *Arabidopsis thaliana*, *Oryza sativa* and *Cucumis sativus* were aligned using MEGA X v10.2 (https://www.megasoftware.net, accessed on 02 June, 2025) (Kumar et al., 2018). Multiple sequence alignment was performed prior to constructing a maximum-likelihood (ML) phylogenetic tree with 1,000 bootstrap replicates. The resulting tree was refined and visualized using iTOL v7 (https://itol.embl.de/, accessed on 02 June, 2025). A combined phylogenetic tree was generated for systematic classification of *CsSNARE* genes, with gene nomenclature assigned according to evolutionary relationships. Functional annotations for identified *CsSNARE* genes were extracted from the genome reference GFF3 file. Chromosomal coordinates of *CsSNARE* sequences were mapped using TBtools v2.154 (https://github.com/CJ-Chen/TBtools/releases, accessed on 02 June, 2025) (Chen et al., 2023) to generate a physical location map.

### 2.3 Conserved motifs, promoter analyses, and gene structure analysis of CsSNARE

Conserved motifs of CsSNARE proteins were identified using the Multiple Expectation Maximization for Motif Elicitation (MEME) suite (https://meme-suite.org/meme/, accessed on 03 June, 2025). Gene structures comprising intron-exon organization were characterized via the Gene Structure Display Server (http://gsds.cbi.pku.edu.cn/, accessed on 03 June, 2025) (Hu et al., 2015). For promoter analysis, 2,000 bp upstream sequences flanking cucumber *CsSNARE* genes were submitted to PlantCARE (https://bioinformatics.psb.ugent.be/webtools/plantcare, accessed on 03 June, 2025) (Lescot et al., 2002) to investigate cis-acting elements.

### 2.4 Duplication and covariance analysis of CsSNARE genes

*CsSNARE* gene duplication events were analyzed using the MCScanX of TBtools, with the criterion of long genes covering 75% of the comparable sequence and 75% of the similarity comparison region in length. Covariance analysis was performed using TBtools among cucumber, *Oryza sativa*, and Arabidopsis, respectively.

### 2.5 In silico expression profiling of CsSNARE genes

To investigate the tissue-specific and developmental expression profiles of *CsSNARE* genes in cucumber, as well as their responsiveness to diverse abiotic (cold, heat, salt) and biotic stresses (Angular leaf spot, Gray mold, Powdery mildew, *Cladosporium cucumerinum*), we retrieved relevant expression datasets from the Cucumber-DB database (http://www.cucumberdb.com/, accessed on 07 June, 2025). Subsequent expression profiling analysis was performed using TBtools software.

### 2.6 Plant materials, growth conditions, and treatments

Cucumber seeds (commercial cultivar 9930) were acquired from Weimibio Co., Ltd. and utilized exclusively for research purposes. The *CsSYP121* gene (*CsaV4_6G002929*) was amplified from cucumber leaf tissue and subsequently cloned into a plant transformation vector (driven by the CaMV 35S promoter, vector acquired from Weimibio Co., Ltd.). The result construct was verified by Sanger sequencing (Sangon Biotech). The recombinant plasmid was introduced into *A. tumefaciens* strain EHA105 via freeze-thaw transformation. Agrobacterium-mediated cucumber transformation was performed by Weimibio Co., Ltd. (http://www.wimibio.com), following established protocols. Briefly, cotyledon explants from 2-day-old germinated cucumber line 9930 seedlings were incubated with 50 mL of recombinant *A. tumefaciens* (OD_600_=0.6) suspension for 30 minutes at 28°C. Following this, explants underwent sequential co-culture phases: an initial 3-day dark incubation at 23 °C followed by 7 days under illumination at 25 °C. The light source was cool white fluorescent tubes (Philips TL-D 36W/33-640, color temperature 4,100 K, spectrum 400–700 nm), providing a photosynthetic photon flux density (PPFD) of approximately 71 μmol·m^-2^·s^-1^, with a photoperiod of 16 h light/8 h dark. Infected tissues were then washed and transferred to shoot induction medium (SIM) supplemented with 50 mg/L spectinomycin, with explants subcultured onto fresh SIM every two weeks during the six-week selection period. Adventitious shoot meristems typically emerged from the cut edges of cotyledon explants after 2–3 weeks of culture on SIM. Surviving explants were subsequently transferred to elongation and rooting media containing the same antibiotic concentration. All tissue culture procedures were performed under strictly sterile conditions in laminar flow hoods, with cultures maintained in sterile vessels (Magenta boxes or parafilm-sealed Petri dishes); non-sterile greenhouse conditions were introduced only after regenerated plantlets with well-developed root systems were successfully acclimatized. Putative transgenic lines, confirmed by PCR analysis using *aadA* gene-specific flanking primers, were acclimatized in greenhouse conditions for seed production.

The *CsSYP121* over-expressing (OE) seeds were germinated in a tray containing vermiculite and nutrient soil at a ratio of 1:1 and cultivated in the growth chamber under the following conditions: temperature of 25-30℃, a light intensity of 250 μmol·m⁻²·s⁻¹, a photoperiod of 16 h (day)/8 h (night), and a relative humidity of 40-50%. When the cucumber seedlings reached three leaves and one heart stage (three fully expanded cotyledons and visible apical meristem), the plants with consistent growth condition were selected for drought stress. Plants were randomly allocated to two treatment groups: (1) well-watered control; (2) drought-stressed treatment subjected to complete water withholding for 14 days; (3) salt-stressed treatment subjected to 150 mmol/L NaCl for 10 days (Zhou et al., 2022). Each treatment comprised three biological replicates with ten plants per replicate. Leaf tissue samples were collected at predetermined time points from the second fully expanded leaf. Harvested samples were immediately frozen in liquid nitrogen and stored at -80°C prior to downstream analysis.

### 2.7 RNA extraction and qRT-PCR

The RNA extraction and qRT-PCR were performed as described before (Fan et al., 2024; Guo et al., 2024; Liang et al., 2024). Total RNA was extracted using TransZol™UP Plus RNA Kit (TransGen Biotech, Beijing, China). The first-strand cDNA was synthesized using EasyScript® One-Step gDNA Removal and cDNA Synthesis SuperMix Kit (TransGen Biotech, Beijing, China) according to the manufacturer’s instructions. The primers of *CsSNARE* genes were synthesized by Shanghai Sangon Biotech (As shown in supplement Table 1). *CsEF1α* (*CsaV4_5G001996*) and *CsCACS* (*CsaV4_3G004932*) were used as internal references. Quantitative real-time PCR (qRT-PCR) was performed using PerfectStart® Green qPCR SuperMix Kit (TransGen Biotech, Beijing, China). Each experiment included three technical replicates and three biological replicates to ensure the reliability of the results.

### 2.8 Western blot

To verify protein expression, plant tissue extracts were subjected to immunoblot (Western blot) analysis using an anti-CsSYP121 antibody, as previously described (Zhang et al., 2018).

### 2.9 Analysis of cation contents, H_2_O_2_ accumulation, and antioxidant enzyme activities

The leaf samples were collected as described above. The analysis of cation content was conducted by Qidian Chemical Technology Service (Liaocheng, China) as previous described (Chen et al., 2016; Grefen et al., 2015; Guo et al., 2024; Liu et al., 2023). After harvesting, the samples underwent acid digestion. The concentrations of Na^+^ and K^+^ ions were determined using inductively coupled plasma optical emission spectrometry (ICP-OES 730, Agilent, USA).

The determination of indicators related to ROS was conducted using commercial kits from Suzhou Keming Biotechnology Co. ltd (China), aligning with prior research (Guo et al., 2024). Fresh leaf tissue (0.1 g) was homogenized in 1 mL of the specific extraction buffer provided with each kit on ice, followed by centrifugation at 12,000 × g for 15 min at 4°C. The resulting supernatant was used for subsequent assays. Specifically, the contents of hydrogen peroxide (H_2_O_2_, H_2_O_2_-2-Y), along with the activities of catalase (CAT, CAT-2-Y), superoxide dismutase (SOD, SOD-2-Y), and peroxidase (POD, POD-2-Y) were assessed. Peroxidase (POD) Activity was determined using the POD-2-Y kit. The assay measures the oxidation of guaiacol in the presence of H_2_O_2_. The reaction mixture contained 50 mM phosphate buffer (pH 7.0), 20 mM guaiacol, 10 mM H_2_O_2_, and enzyme extract. The increase in absorbance at 470 nm due to tetraguaiacol formation (ε_470_= 26.6 mM^-1^cm^-1^) was recorded. One unit of POD activity was defined as an absorbance change of 0.01 per minute per mg protein. This method primarily detects the activity of class III (guaiacol-type) peroxidases.

### 2.10 Detection of protein subcellular localization

The template cDNA for the cloning of CsSYP121 was derived from the aforementioned reverse transcription. Amplification primers were designed to span the coding sequence (CDS) region of the target gene, excluding the promoter sequence, with the objective of obtaining the open reading frame (ORF) for subsequent Agrobacterium tumefaciens-mediated transient expression assays in tobacco. High-fidelity DNA polymerase KOD FX was employed to ensure amplification accuracy, and all primers were synthesized by Shanghai Sangon Biotechnology Co., Ltd. (detailed in Supplementary Table 1). Gateway® attB1 and attB2 recombination sites were incorporated into the amplicons. Entry clones were constructed via BP recombination using the pDONR207 vector (Invitrogen), and the final expression clones were generated through LR recombination utilizing LR Clonase™ II enzyme mix (Invitrogen).

To investigate the protein localization, *CsSYP121* was cloned into the pUBN-GFP-DEST vector via LR reaction, following established protocols (Grefen et al., 2010b; Zhang et al., 2015). And AtAHA1 was cloned into the pUBC-YFP-DEST vector as a marker via the same LR reaction. *Agrobacterium tumefaciens* GV3101 harboring this construct was used to infiltrate tobacco (*Nicotiana tabacum*) leaves, as previously described (Zhang et al., 2015). GFP fluorescence was observed and imaged using a confocal laser scanning microscope at 48-72 h post-transfection. For the GFP localization analysis, confocal microscopy was performed using a Leica STELLARIS 5 equipped with a 20× objective. The GFP signal was excited with a 488 nm laser and detected through a 505-530 nm bandpass filter.

### 2.11 Statistical analysis

The sample size (n) for each comparison refers to the number of biological replicates. Statistical analysis of independent experiments was reported as means±SE as appropriate, with signifcance determined by Student’s t-test or ANOVA.

## 3. Results

### 3.1 Identification of CsSNARE genes in cucumber

Based on the sequence of homology with Arabidopsis, we identified 94 putative *CsSNARE* gene candidates in cucumber. Through analysis of protein length and structural integrity, 51 members of the *CsSNARE* gene family were finally obtained and named according to their affinities to homologous of Arabidopsis and *O. sativa* (Table 1). Bioinformatic analysis revealed substantial diversity in molecular mass (10.9-37.5 kDa) across the CsSNARE family, except for the Tomosyn-like R-CsSNARE protein CsTYN11 (CsaV4_5G000834) with 121.5 kDa and CsTYN12 (CsaV4_3G000037) with 116.6 kDa, which were above the range. To investigate evolutionary dynamics, a comprehensive phylogenetic analysis was performed integrating *CsSNARE* sequences with their Arabidopsis and *O. sativa* counterparts (Figure 1). This analysis resolved the *CsSNARE* family into five discrete clades: *Qa-CsSNARE* (14 members), *Qb-CsSNARE* (9 members), *Qc-CsSNARE* (10 members), *Qb+c-CsSNARE* (3 members), and *R-CsSNARE* (15 members). The classification scheme aligns with established SNARE complex composition principles (Zhang et al., 2020), providing a framework for functional characterization of these key regulators in vesicle trafficking.

**Fig. 1.**
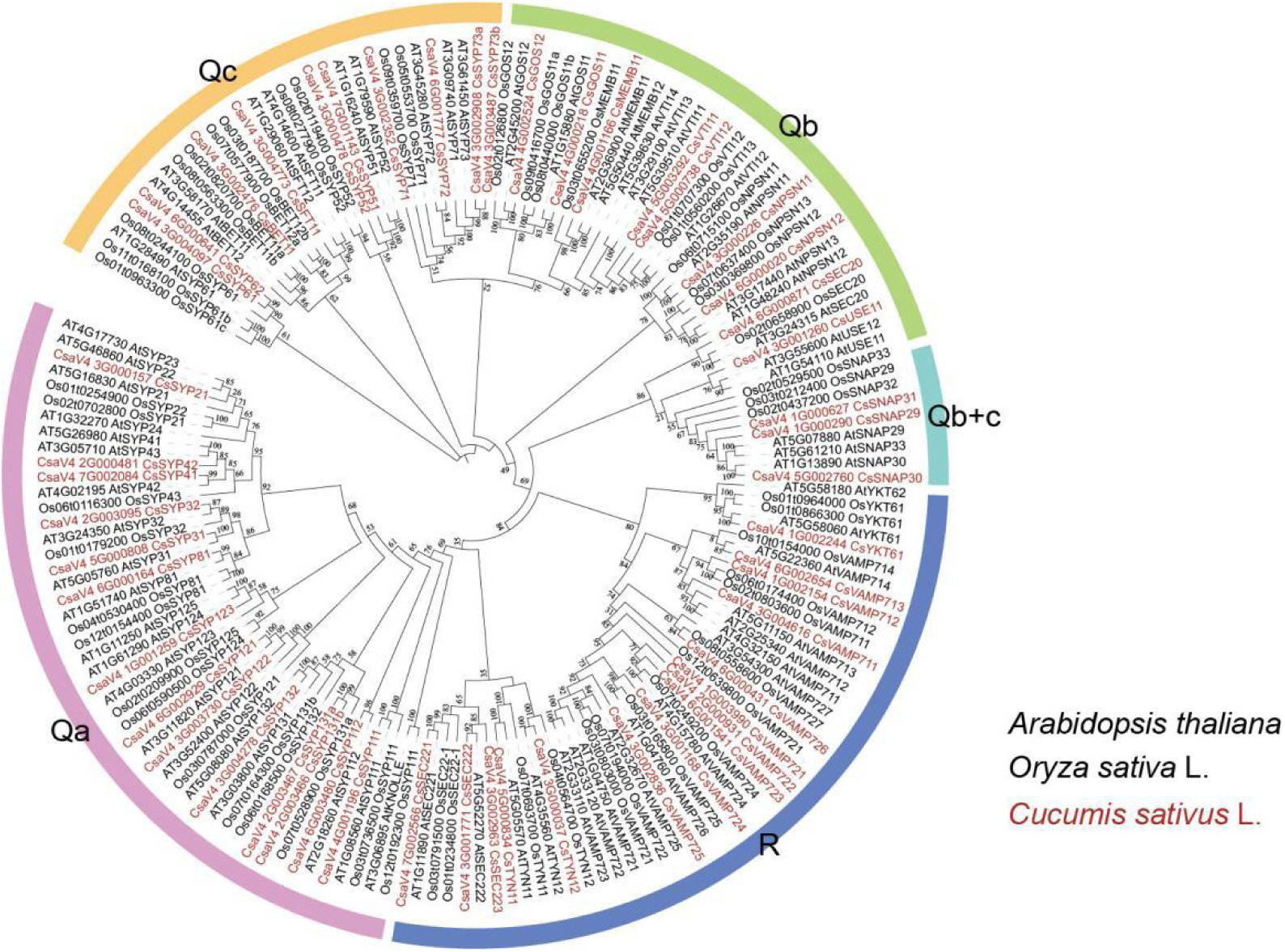
Phylogenetic analysis of 51 *CsSNARE* genes identified in Cucumber. Different colours indicate different SNARE classifications.

**Table 1.**
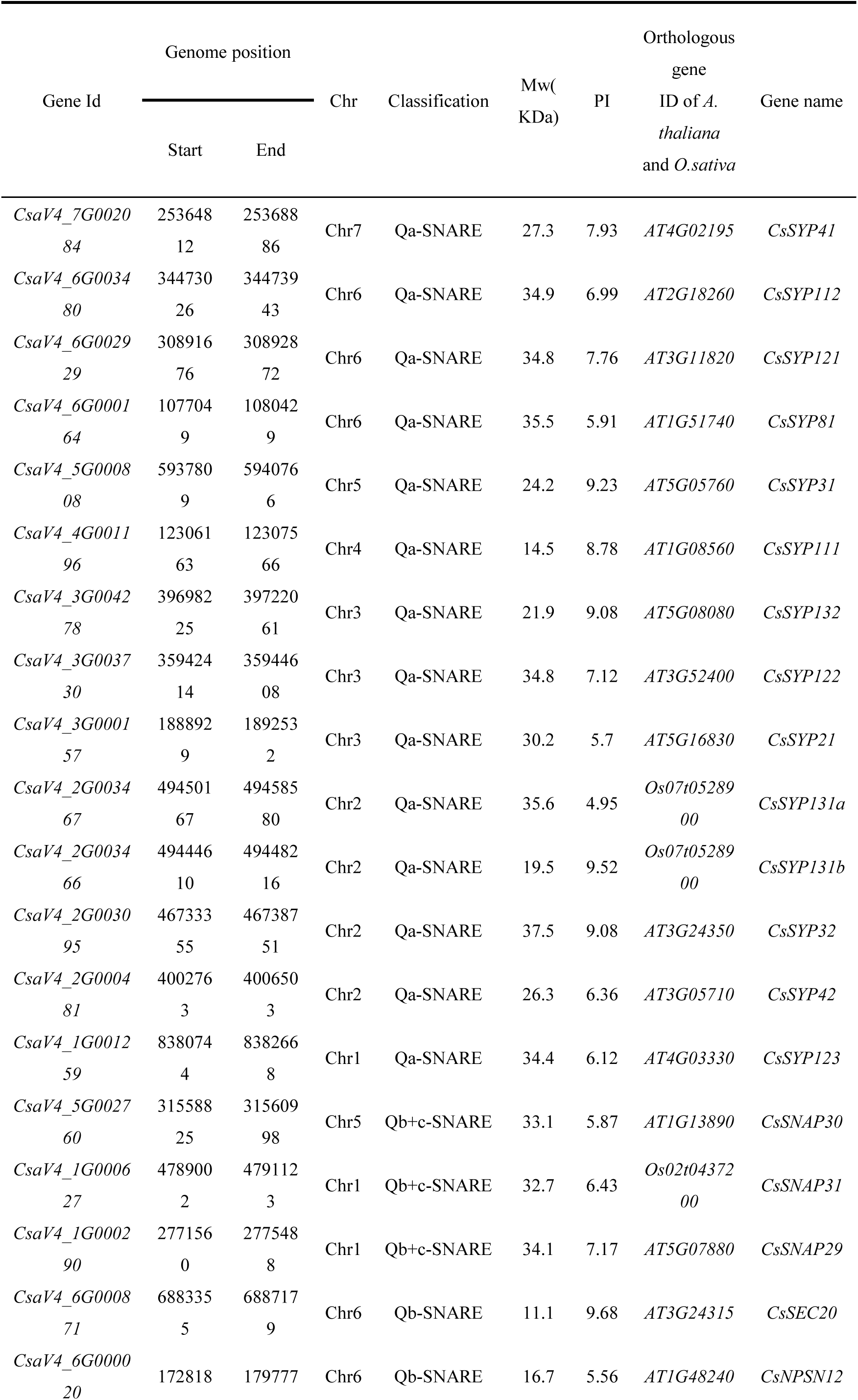
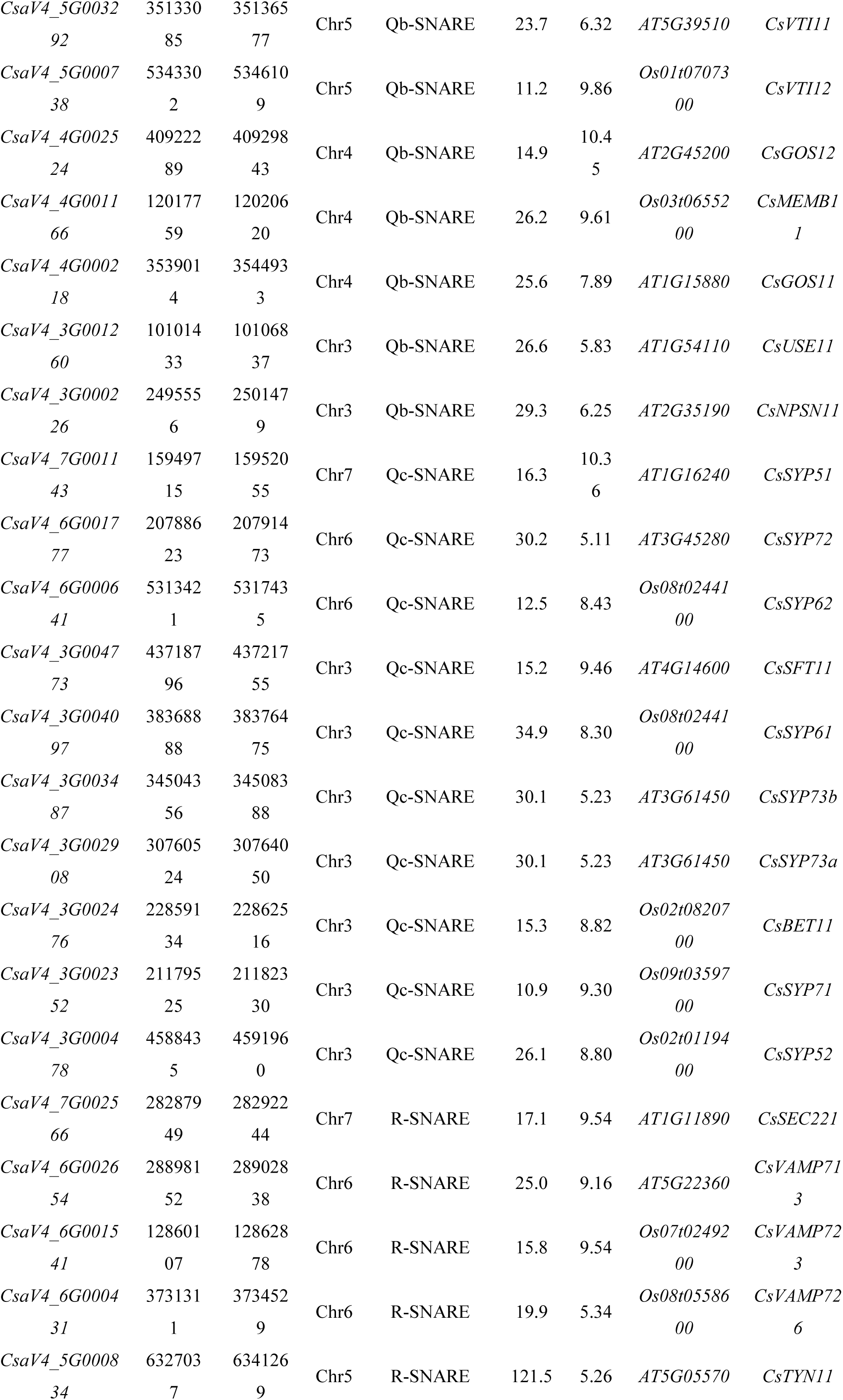
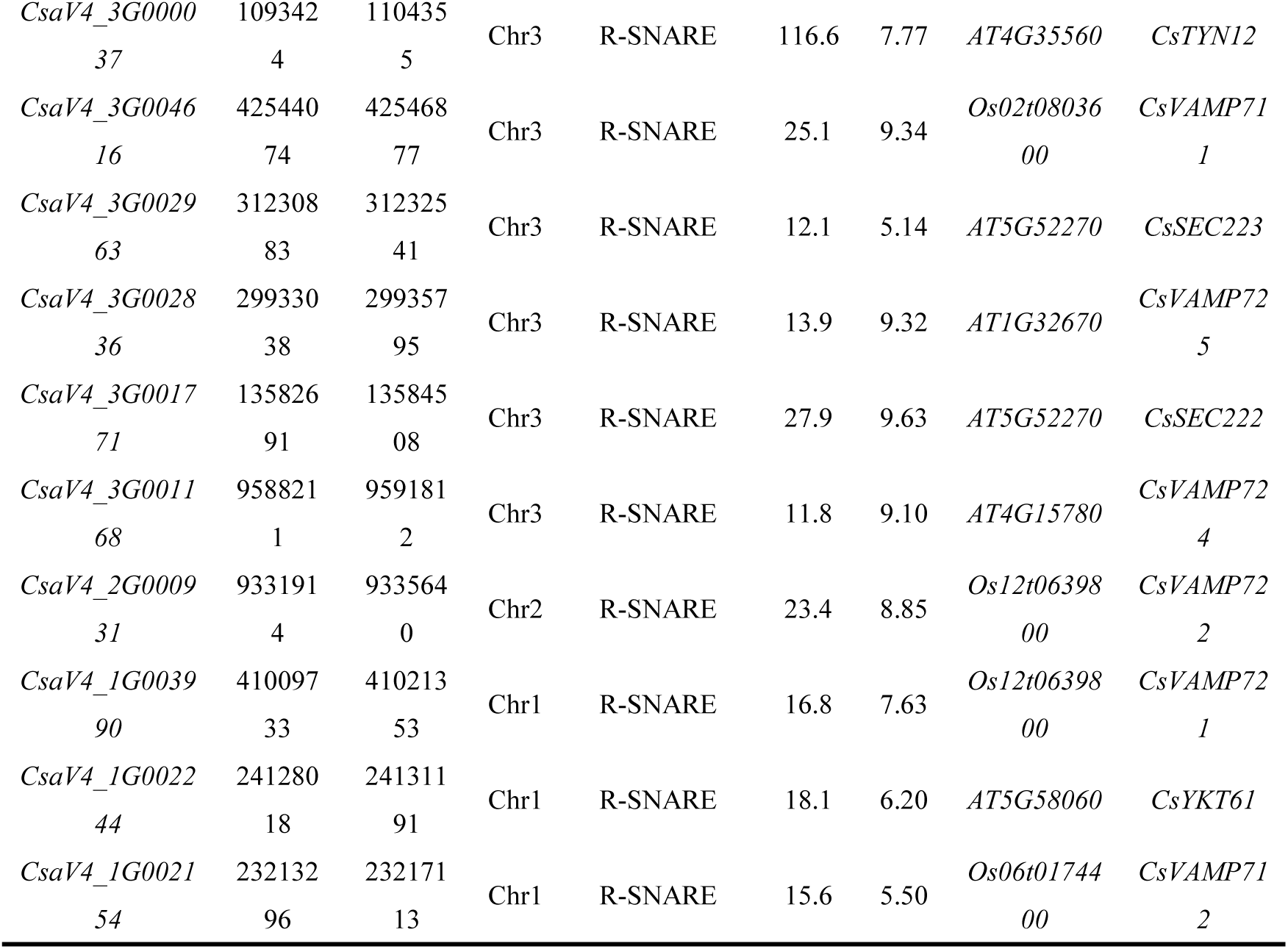
Detailed information for *CsSNARE* genes in *C. sativus* L.

### 3.2 Intrachromosomal distribution, exon-intron structure, conserved protein motifs, and cis-acting elements in the promoter region of CsSNARE genes

Figure 2 illustrates the non-random distribution pattern of *CsSNARE* genes across seven cucumber chromosomes. Notably, chromosome 3 emerged as the primary cluster with 18 *CsSNARE* genes, followed by chromosome 6 (10 genes) and chromosome 1 (6 genes). Chromosome 2 and 5 each harbored 5 *CsSNARE* genes. Chromosomes 4 and 7 each harbored 4 and 3 *CsSNARE* genes. This heterogeneous chromosomal localization suggests potential functional differentiation among *CsSNARE* genes.

**Fig. 2.**
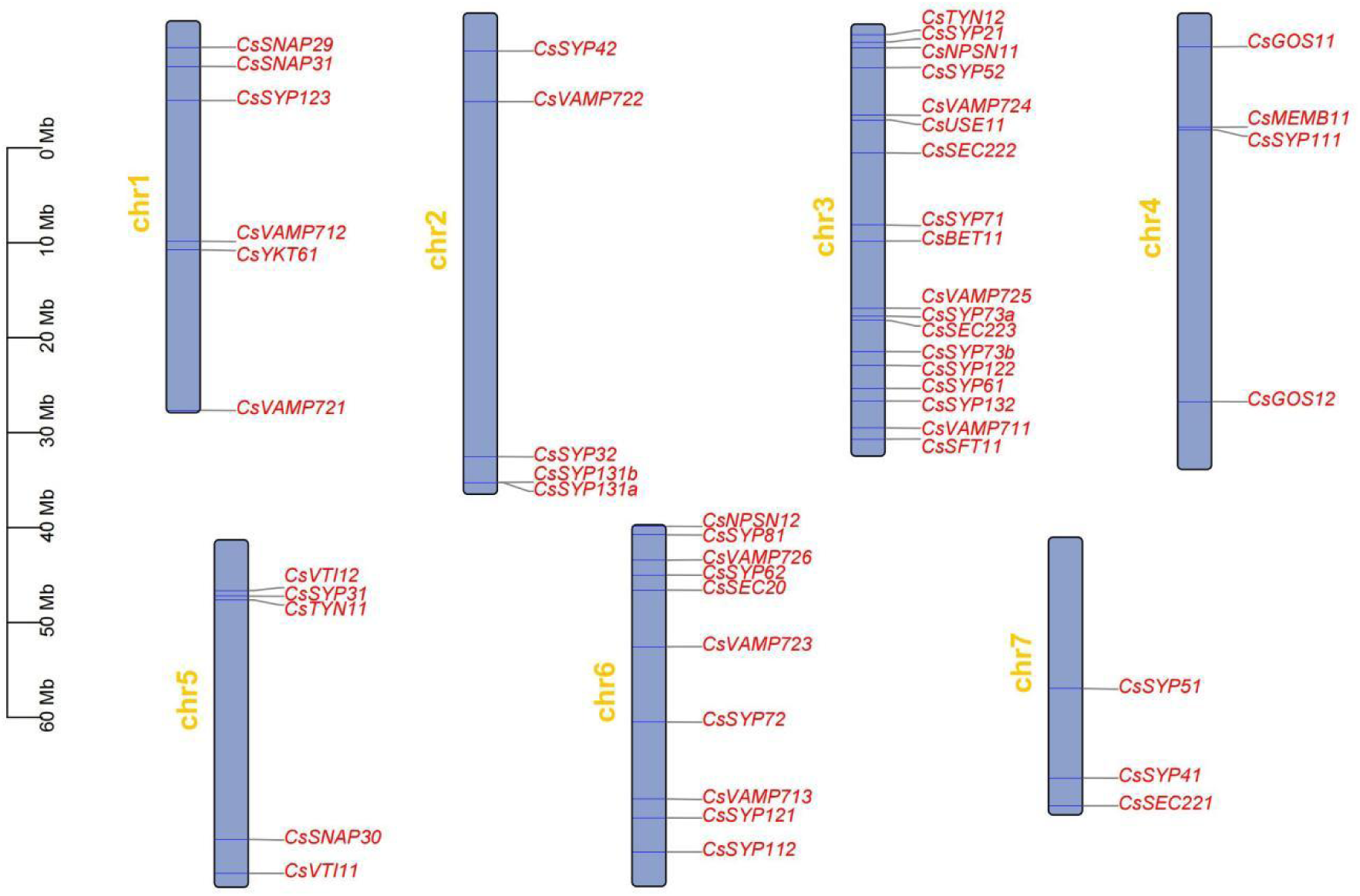
Chromosomal distribution of *CsSNARE* genes in cucumber. The chromosomal locations of 51 *CsSNARE* genes were retrieved from the Cucumber Multi-omics Database (CLv4.0)

The exon-intron structures of *CsSNARE* genes are shown on the right side of Fig. 3. Except for *CsSYP21*, *CsSYP42*, *CsSYP81*, *CsSYP112*, *CsGOS11*, *CsSFT11* and *CsSYP61* which only contain coding sequences (CDS), other *CsSNARE* members all contain untranslated regions (UTR) and CDS. Members with closer kinship contain similar genetic structures.

**Fig. 3.**
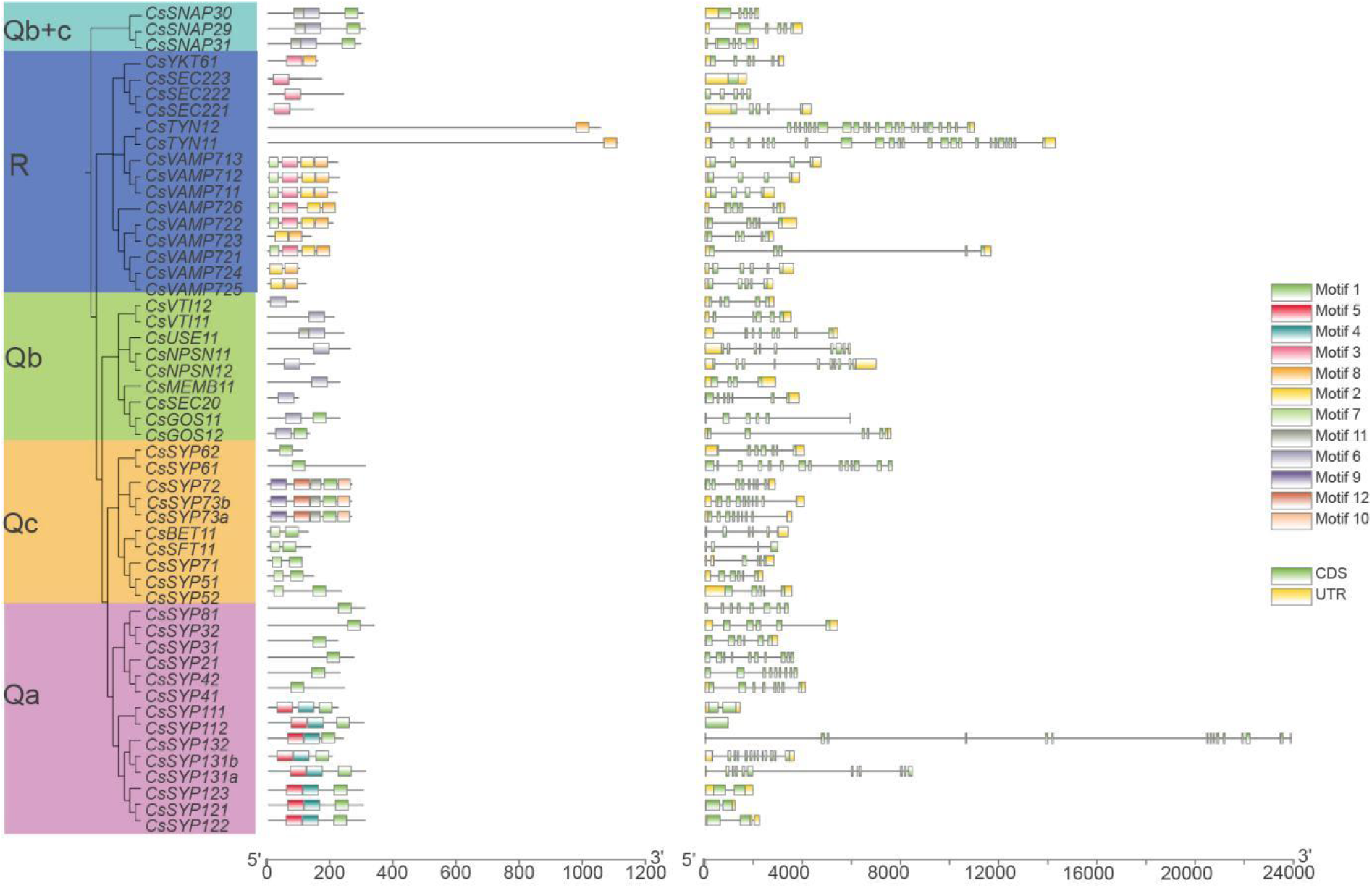
Protein motifs and gene structures of *CsSNARE* from Cucumber. (Left) In the CsSNARE proteins, 12 motifs were identified by the MEME tool, represented by different colors (1-12) and depicted by TBtools. (Right) The exon-intron structure of these *CsSNARE* genes were predicted by TBtools. The yellow boxes represent the coding sequence (CDS), the black lines represent introns, and the green boxes represent up/downstream untranslated regions (UTR).

**Fig. 4.**
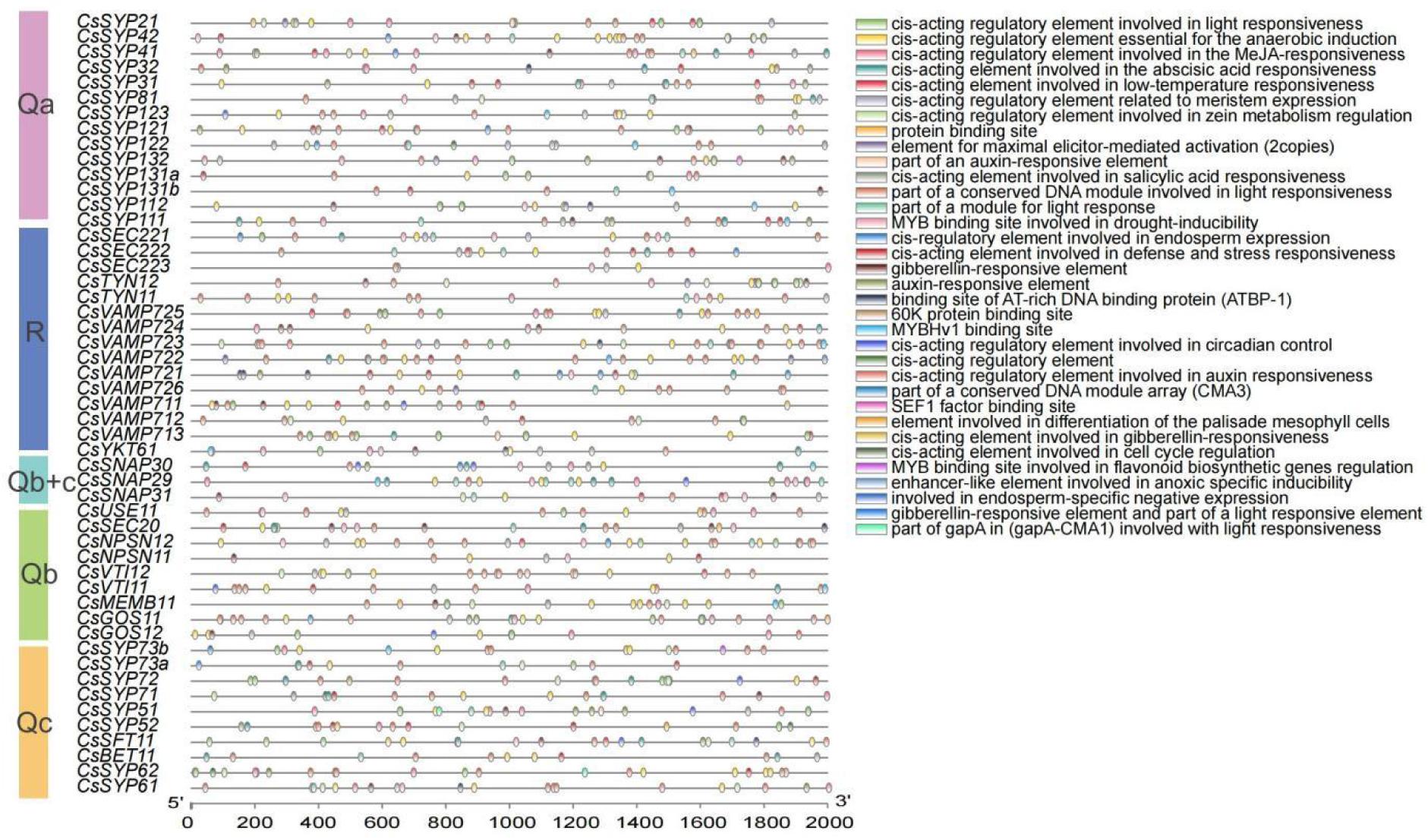
*Cis*-acting element analysis of the promoter regions of *CsSNARE* genes in Cucumber. Based on the functional annotations, the cisacting elements were shown in boxes with different colors.

Conserved motif characterization of CsSNARE proteins was conducted via the MEME suite, revealing 12 distinct conserved motifs across the family. As depicted in Figure 3 (left panel), motif distribution exhibited subfamily-specific patterns: R-CsSNARE members predominantly featured Motifs 2, 3, 4, 7, and 8; Qa-CsSNAREs uniquely harbored Motif 5; Qc-CsSNAREs co-expressed Motifs 1, 7, 8, 9, 10, 11, and 12; Qb-CsSNAREs contained Motifs 1, 6, and 11; while Qb+c-CsSNAREs exclusively retained Motif 1, 6, and 11. Notably, Except for R-CsSNARE, motif 1 is widespread in other subfamilies, implicating its potential role as a fundamental component of the CsSNARE core structure. Motif 5 localized to the N-terminal Habc domain in most Qa-CsSNAREs. The R-CsSNARE subgroup specifically enriched for Motifs 3 and 7, corresponding to the Longin domain (Motifs 2+8). Similar conserved motifs within the same subfamily indicate that they may have similar functions, while the different distributions of conserved motifs in different subfamilies may be one of the reasons for their functional diversity during the evolutionary process. These motif architectures corroborate the phylogenetic classification of CsSNARE proteins, reinforcing functional distinctions among subfamily members.

To understand how *CsSNARE* genes are regulated, we analyzed cis-acting elements (CREs) in the 2,000 bp region upstream of each gene. We identified 34 types of CREs and grouped them into three main categories. Firstly, Light-Responsive Elements, including part of a conserved DNA module involved in light responsiveness (182 occurrences, 18.5%), *cis*-acting regulatory element involved in light responsiveness (115 occurrences, 11.7%), and part of a module for light response (28 occurrences, 2.8%). Second, Plant Hormone-Responsive Elements, including *cis*-acting regulatory element involved in MeJA-responsiveness (120 occurrences, 12.2%), *cis*-acting element involved in abscisic acid responsiveness (103 occurrences, 10.5%), *cis*-acting element involved in salicylic acid responsiveness (33 occurrences, 3.4%), Gibberellin-responsive element (28 occurrences, 2.8%), Auxin-responsive element (24 occurrences, 2.4%). Third, Stress-Responsive Elements, including *cis*-acting regulatory element essential for anaerobic induction (117 occurrences, 11.9%), *cis*-acting element involved in defense and stress responsiveness (34 occurrences, 3.5%), MYB binding site involved in drought-inducibility (29 occurrences, 3.0%), *cis*-acting element involved in low-temperature responsiveness (15 occurrences, 1.5%). Additionally, we found Auxiliary Regulatory Elements, including *cis*-acting regulatory element involved in circadian control (10 occurrences), *cis*-acting regulatory element involved in zein metabolism regulation (33 occurrences), *cis*-acting regulatory element related to meristem expression (18 occurrences), and CRE involved in endosperm expression (10 occurrences). The abundance of stress-related and hormone-responsive motifs suggests the CsSNARE family helps plants respond to various stresses and hormones.

### 3.3 Intra- and inter-species covariance analysis of CsSNARE genes in cucumber

Next, we analyzed the evolutionary dynamics of *CsSNARE* gene duplication and identified conserved genomic regions across species. Through within-species comparative analysis of cucumber *CsSNARE* genes and interspecific synteny assessments with Arabidopsis and rice *SNARE* gene sets, key insights into genomic evolution were obtained. Figure 5A shows the chromosomal distribution of *CsSNARE* subfamilies (*CsSYPs*, *CsVAMPs*, *CsSNAPs*). This distribution suggests gene duplication drove family expansion. We identified seven segmental duplication events in cucumber. Chr3 was a hotspot, containing five events. Collinear gene pairs indicate possible functional similarities. Examples include: *CsVAMP724* on chromosome 3 is collinear with *CsVAMP725* on chromosome 3 and *CsVAMP722* on chromosome 2, and *CsSNAP30* on chromosome 5 is collinear with *CsSNAP29* and *CsSNAP31* on chromosome 1. It indicates that the *CsSNARE* gene may have undergone gene replication or gene drift, resulting in possible similarities in its function and expression pattern. This pattern implies segmental duplicative expansion as a major evolutionary force shaping the *CsSNARE* gene family.

**Fig. 5.**
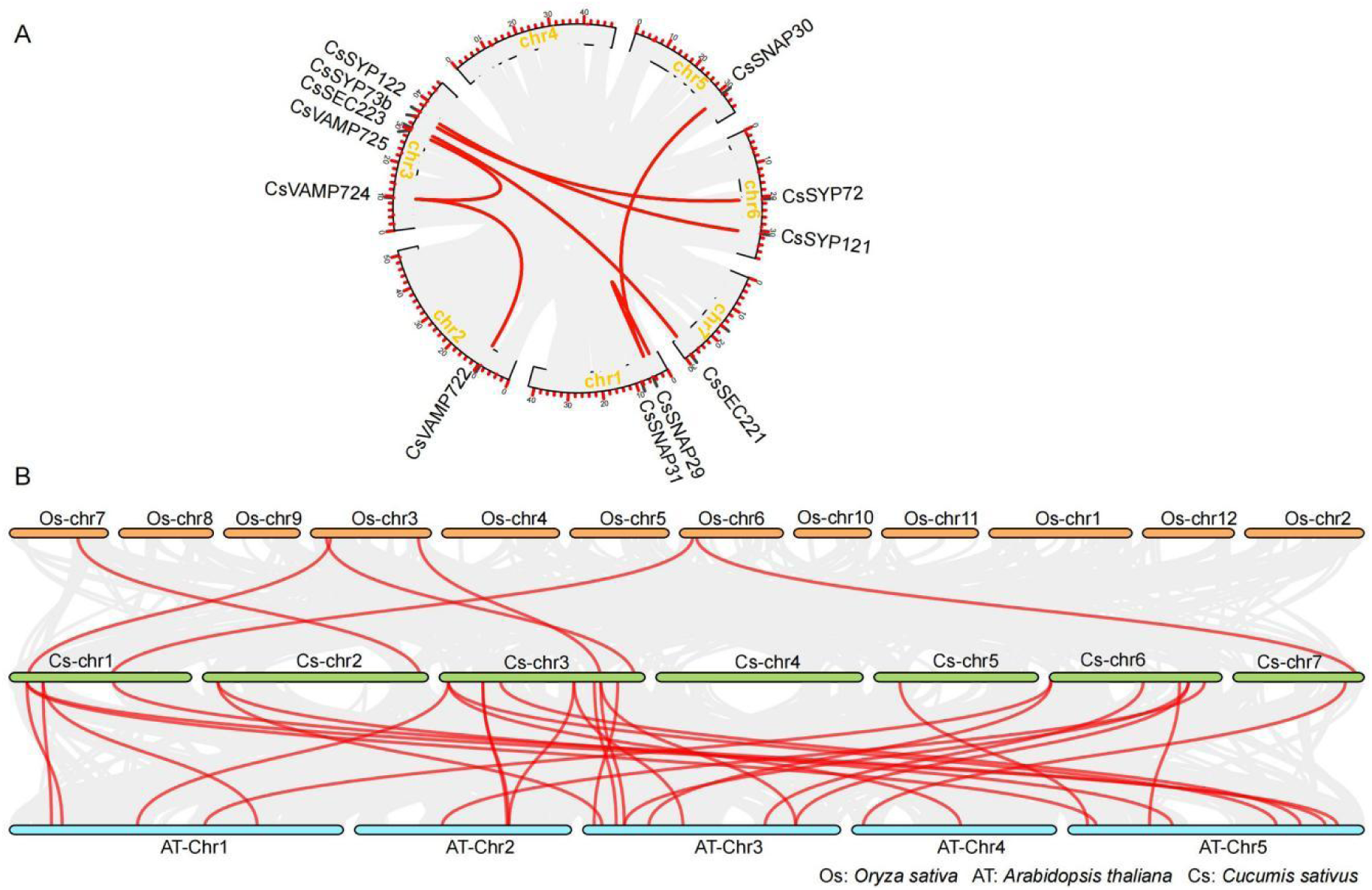
Intra- and inter-species covariance analysis of the *CsSNARE* gene family in Cucumber. (A) Collinearity analysis within the *CsSNARE* genes. (B) Collinearity analysis among *SNARE* genes. AT stands for *A. thaliana*, Cs for *C. sativus*, and Os for *O. sativa*. The *SNARE* gene pairs replicated in fragments are linked through red lines between chromosomes.

Synteny analysis revealed stronger conservation between cucumber and Arabidopsis (29 orthologous pairs) than between cucumber and rice (6 pairs). This reflects closer evolutionary relatedness between these dicots than between cucumber (dicot) and rice (monocot). These findings suggest distinct evolutionary paths shaped by lineage-specific duplications and selection.

### 3.4 In silico transcript analysis of CsSNARE genes in cucumber

To understand *CsSNARE* functions, we analyzed their expression across tissues, organs, and developmental stages using public RNA-seq data. Figure 6 shows ubiquitous *CsSNARE* expression. Transcripts accumulated most in roots, hypocotyls, tendrils, and stems, suggesting key roles in vegetative growth and development. Within the *Qa-CsSNARE* subfamily, *CsSYP123* demonstrated remarkable upregulation in hypocotyl tissue, implicating its potential involvement in seedling establishment and early growth regulation. Concurrently, *CsSYP121* displayed significant enrichment in primary root tissues, potentially mediating fundamental processes such as ion uptake, nutrient assimilation, and osmotic stress responses. The apical meristem-specific induction of *CsSYP112*, *CsSYP41* and *CsSYP111* further supports its postulated function in governing morphogenic programs and axial patterning during shoot development. Regarding leaf-specific expression, *Qc-CsSNARE* member *CsSYP61* consistently exhibited elevated transcript abundance throughout leaf maturation stages, indicating stage-specific regulatory functions potentially associated with foliar morphogenesis or photosynthetic capacity development. These tissue- and stage-specific patterns highlight how *CsSNARE* genes coordinate plant growth and adaptation.

**Fig. 6.**
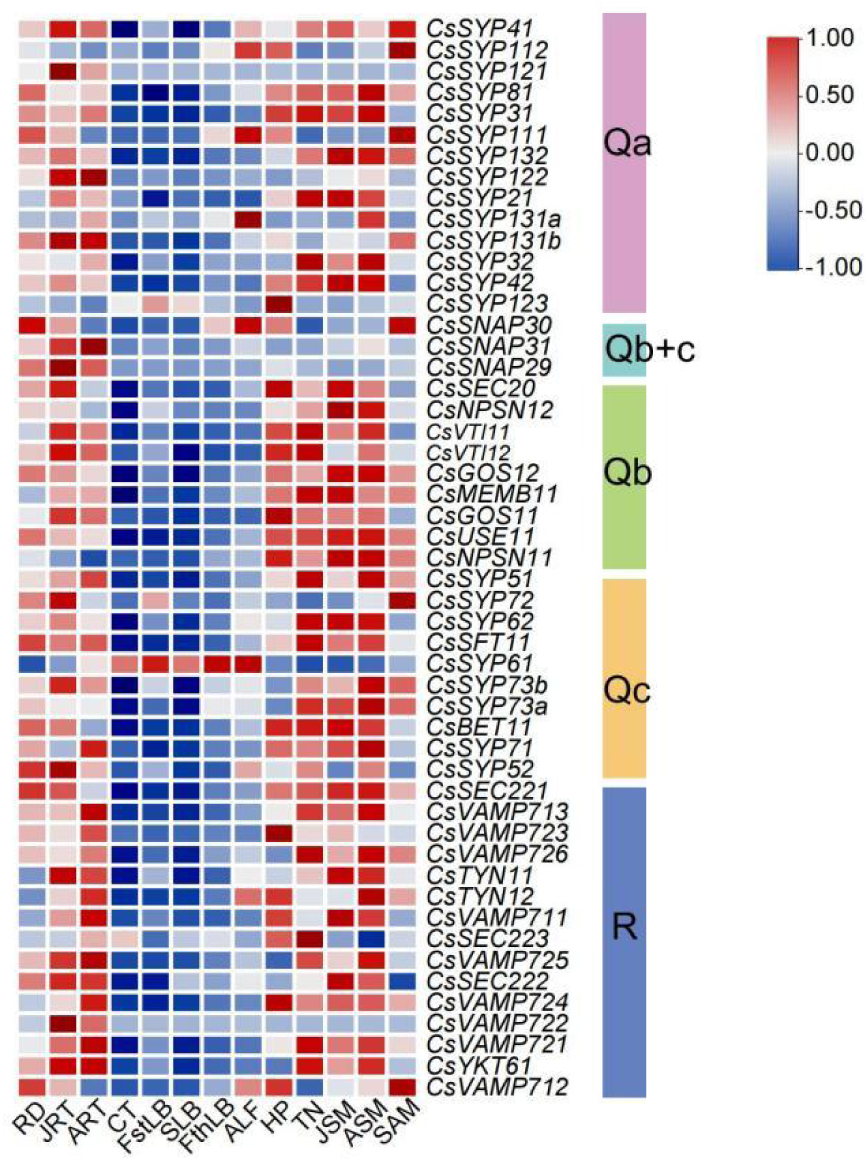
Expression profile of *CsSNARE* genes in different tissues and developmental stages. RD is radicle on juvenile, JRT is juvenile root, ART is Adult root, CT is Cotyledon, FstLB is first leaf blade, SLB is second leaf blade, FthLB is forth leaf blade, ALF is Adult leaf, HP is Hypocotyl, TN is Tendril, JSM is juvenile stem, ASM is Adult stem, SAM is Shoot apical meristem. The color scale represents the level of expression (TPM value), from high (red) to low (blue). Gene expression data are downloaded from the Cucumber Multi-omics database.

To elucidate the functional dynamics of the *CsSNARE* gene family under environmental stresses, this study also retrieved transcriptomic profiling data (as TPM values) from the Cucumber Multi-omics Database. Figure 7 shows distinct expression patterns under different stresses. Most *CsSNARE* genes responded differentially to Angular Leaf Spot (ALS) and *Cladosporium cucumerinum* (CC) infections, indicating defense roles. During Gray Mold (GM), *CsSYP61* (*Qc-SNARE*) was uniquely induced, suggesting specific resistance to *Botrytis cinerea*. Most *R-CsSNARE* genes gradually increased during Powdery Mildew (PM) infection, suggesting sustained defense. Cold stress analysis unveiled complex regulatory patterns: *CsSYP123, CsVAMP721, CsNSPN11, CsNSPN12,* and *CsSNAP30* stayed constitutively high. *CsVAMP721* decreased during prolonged cold, suggesting negative feedback. *CsSYP122* and *CsSYP112* peaked at 24 hours. For heat stress, *CsSYP123* showed a transient peak at 20 minutes post-shock. *Qa-SNARE* genes dominated the heat response. Salt stress responsiveness analysis demonstrated temporally coordinated expression changes. *CsSYP132* rose continuously. Most genes peaked at 96 hours. *CsVAMP723*, *CsGOS11*, and *CsVTI12* responded early (48 hours). *CsSNAP30*, *CsSYP61*, *CsVAMP712*, and *CsUSE11* responded rapidly (within 10 hours).

**Fig. 7.**
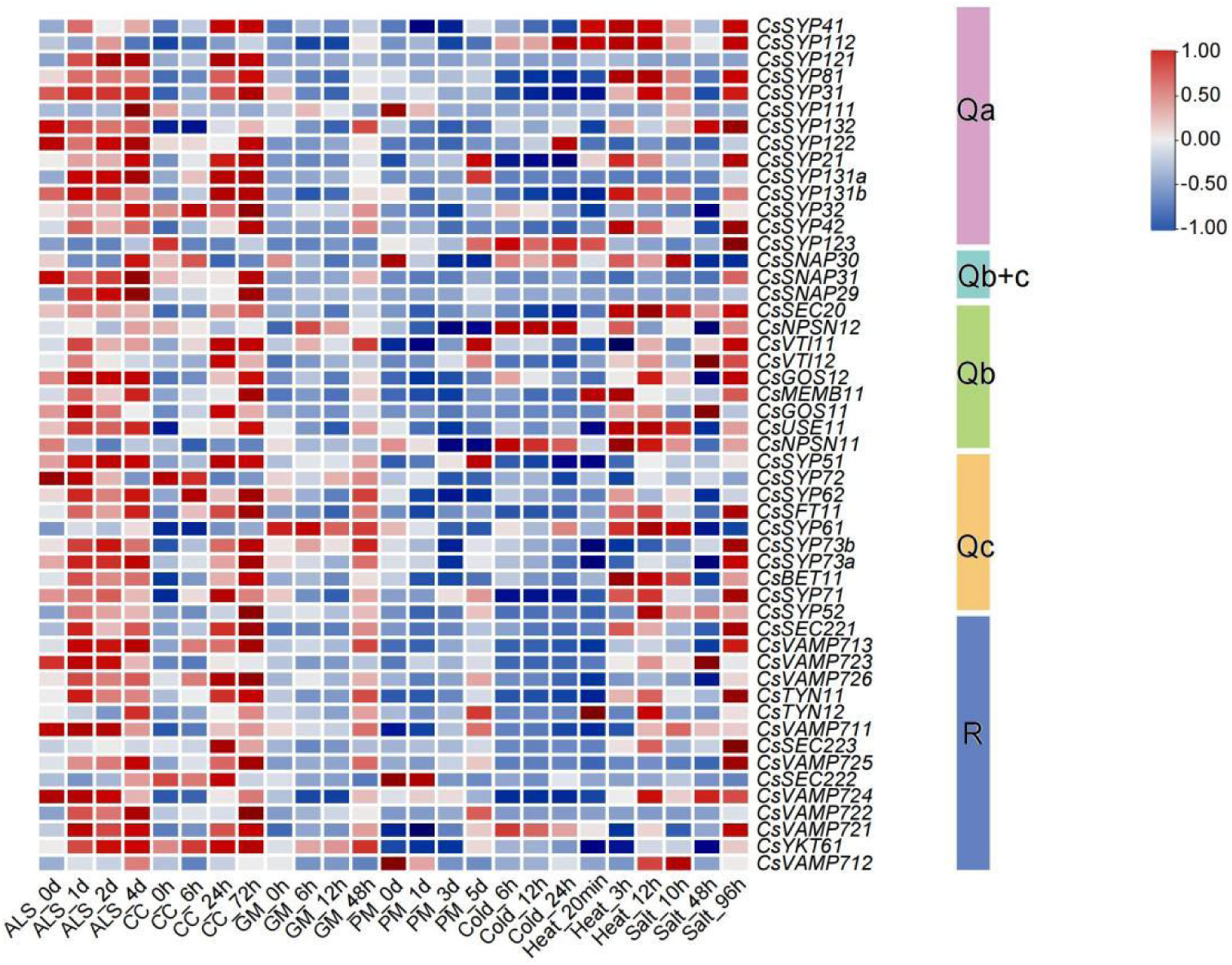
The relative transcript levels of *CsSNARE* genes in Cucumber under different biological and abiotic stresses. ALS is angular leaf spot, CC is *Cladosporium cucumerium*, GM is Gray mold, PM is Powdery mildew. The color scale represents the level of expression (TPM value), from high (red) to low (blue). Gene expression data are downloaded from the Cucumber Multi-omics database.

Collectively, these dynamic expression profiles demonstrate widespread *CsSNARE* involvement in stress adaptation. They likely regulate membrane trafficking, cellular homeostasis, and stress signaling through precise temporal and tissue-specific mechanisms to enhance plant resilience.

### 3.5 Expression profles of CsSNARE genes in response to drought and salt stress

Integration of cis-acting element analyses within cucumber *CsSNARE* promoter regions (Fig. 5) and transcriptomic profiling under stress conditions (Fig. 7) indicated that the majority of *Qa-CsSNARE* genes are transcriptionally responsive to abiotic stresses. Building upon cucumber’s known sensitivity to water deficit and salinity, we prioritized characterizing the drought response of this gene family. To mimic agricultural drought, plants were subjected to a 14-day withholding of irrigation. Subsequent qRT-PCR was employed to systematically resolve the expression dynamics of *Qa-CsSNARE* genes under these controlled conditions. As shown in Figure 8, after 14 days of drought stress, only *CsSYP41* exhibited significant upregulation compared to control plants. In contrast, *CsSYP81*, *CsSYP112*, *CsSYP122*, *CsSYP123*, *CsSYP131b*, and *CsSYP132* displayed significant downregulation at this final timepoint. Notably, the repression of *CsSYP112*, *CsSYP122*, and *CsSYP131b* began significantly earlier, at day 7 of treatment. While *CsSYP42* also showed significant downregulation at day 7, its expression recovered toward control levels by day 14.

**Fig. 8.**
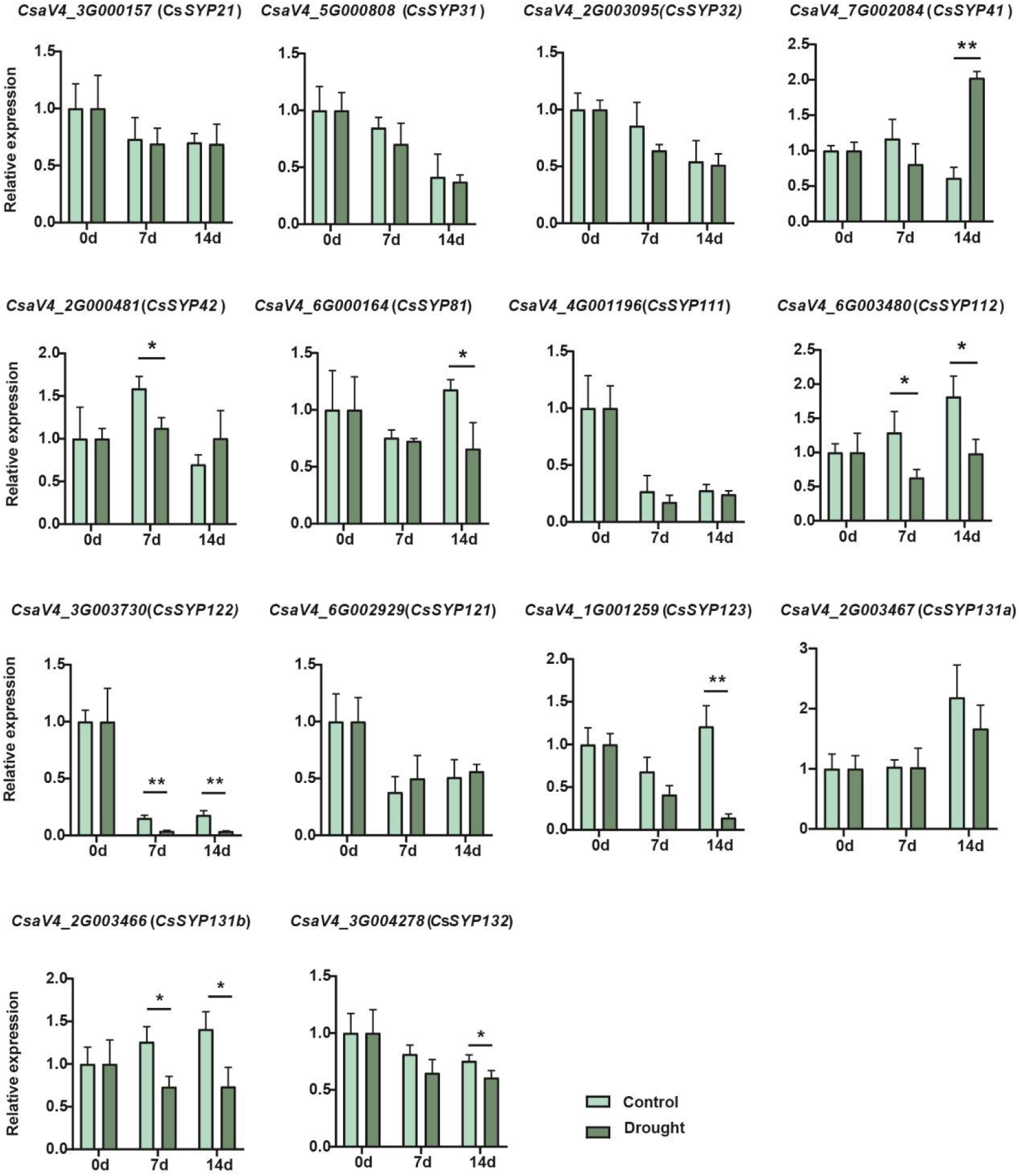
Expression profiles of Qa-CsSNARE genes up-regulated or downregulated in Cucumber under drought stress. The qRT-PCR analyses were used to access transcript levels of *Qa-CsSNARE* genes with or without seven and fourteen days of drought treatment. Each bar represents the mean±SE normalized to *CsEF1α* (*CsaV4_5G001996*) and *CsCACS* (*CsaV4_3G004932*). All samples were run in three biological and three technical replicates. Light green columns represent “Control”, and Dark green columns represent “Drought”. Asterisk indicates that the gene expression under stress has a significant difference compared with the control (\**P*<0.05, \*\**P*<0.01).

To investigate salt stress responses, an experimental group was treated with a 150 mM NaCl solution, whereas the control group received standard cultivation practices. Leaf tissues were collected at 0, 3, and 7 days post-treatment for qRT-PCR analysis. The results revealed significant changes in the expression of *CsSYP21*, *CsSYP31*, *CsSYP32*, *CsSYP111*, *CsSYP122*, *CsSYP121*, *CsSYP123*, and *CsSYP131b* after 7 days of salinity (Fig. 9). Among these, *CsSYP121*, *CsSYP123*, and *CsSYP131b* were strongly induced, whereas the other genes were significantly repressed.

**Fig. 9.**
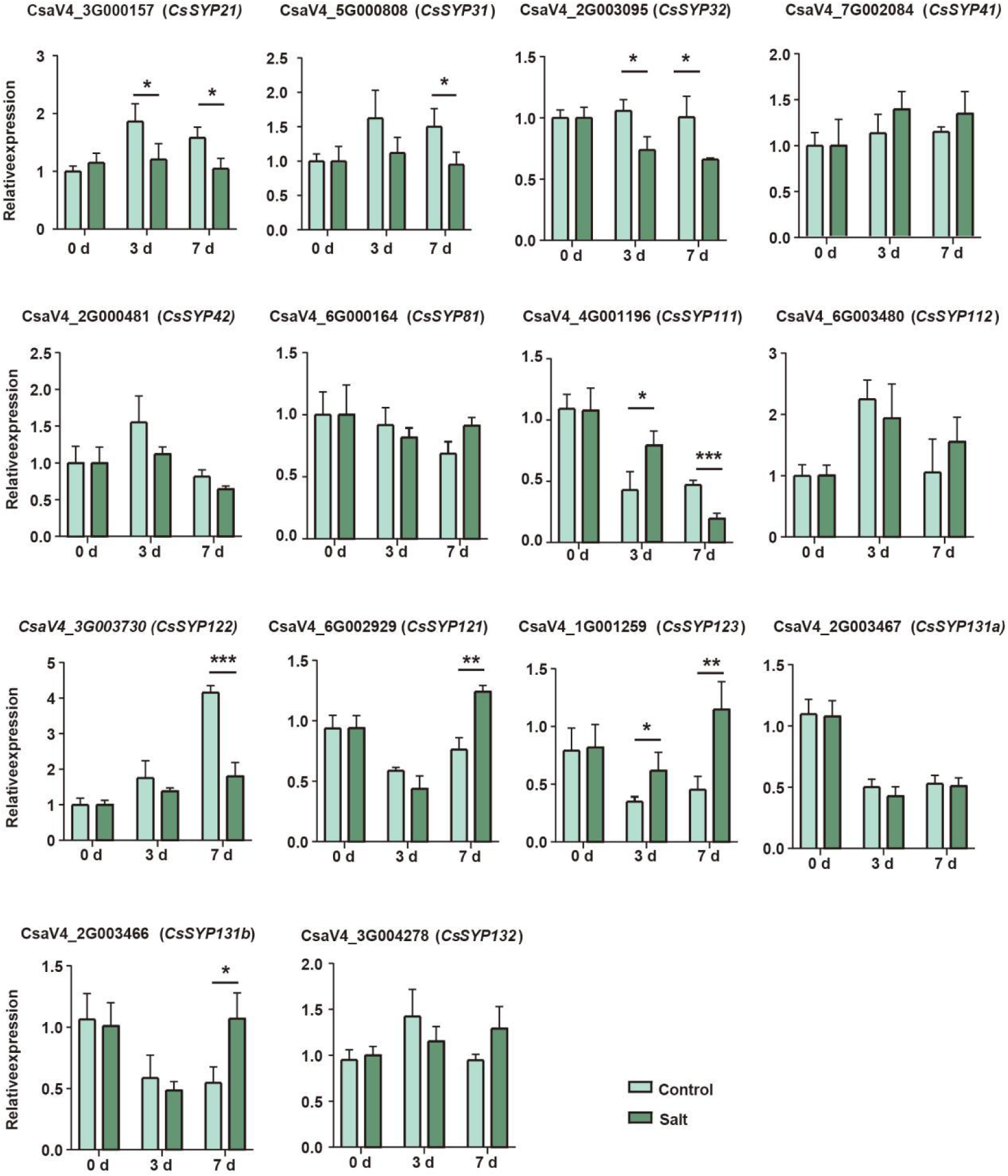
Expression profiles of *Qa-CsSNARE* genes up-regulated or downregulated in wild-type cucumbers under salt stress. The qRT-PCR analyses were used to access transcript levels of *Qa-CsSNARE* genes with or without three and seven days of salt treatment. Each bar represents the mean±SE normalized to *CsEF1α* (*CsaV4_5G001996*) and *CsCACS* (*CsaV4_3G004932*). All samples were run in three biological and three technical replicates. Light green columns represent “Control”, and Dark green columns represent “Salt”. Asterisk indicates that the gene expression after stress has a significant difference compared with the control (\**P*<0.05, \*\**P*<0.01, \*\*\**P*<0.001).

Initial analyses revealed constitutive high expression of *CsSYP121* in roots, as illustrated in Fig. 6. The response of *CsSYP121* expression in cucumber roots to salt stress was evaluated. Fig. S2 demonstrates that *CsSYP121* transcript abundance in roots increased significantly above control levels after 3 days of salt exposure, subsequently returning to baseline levels by day 7. Conversely, a marked and significant upregulation of *CsSYP121* expression was observed in leaves following 7 days of salt treatment. Based on these findings, leaf-related phenotypic changes warrant focused attention in future studies.

### 3.6 Overexpression of CsSYP121 alleviates the inhibitory effects of salt stress on cucumber by maintaining K^+^ and Na^+^ homeostasis and suppressing ROS accumulation

*CsSYP121* is a member of the *Qa-SNARE* family in cucumber. It displays constitutive expression in most cucumber tissues, peaking in the root system (Fig. 6). No substantial changes in its transcript abundance were detected upon exposure to cold, heat, or drought stresses (Fig. 7-9). Moreover, after salt stress, the expression level of *CsSYP121* was significantly higher in both leaves and roots compared to the control group, with a more pronounced increase observed in the leaves (Fig. S2). Given that its ortholog in Arabidopsis and maize modulates plasma membrane K^+^ channel and water channel activities via protein-protein interactions (Honsbein et al. 2009; Besserer et al. 2012; Hachez et al. 2014), we examined the physiological role of *CsSYP121* by overexpressing it in cucumber. The overexpression of *CsSYP121* was confirmed by qRT-PCR (Fig. S3) and western blot (Fig.S4). As shown in Figure 10, under control conditions, *CsSYP121* overexpression had no significant effect on plant height or stem diameter. After 7 days of salt stress treatment, inhibition of plant height and stem diameter was observed. However, *CsSYP121* overexpression alleviated the suppression of cucumber plant height. The 14-day drought stress treatment revealed that the *CsSYP121*-OE lines were as sensitive to drought stress as WT plant (Fig. S1). Combined with the qPCR results in Fig. 8, which showed that CsSYP121 was not significantly induced under drought stress, we speculate that this gene may functionally prioritize the regulation of ionic stress, with a relatively limited response to osmotic stress. Additionally, subcellular localization analysis of the CsSYP121 protein was performed, and the results showed that CsSYP121 co-localizes with the marker protein AtAHA1 at the plasma membrane (Fig. 10 E).

**Fig. 10.**
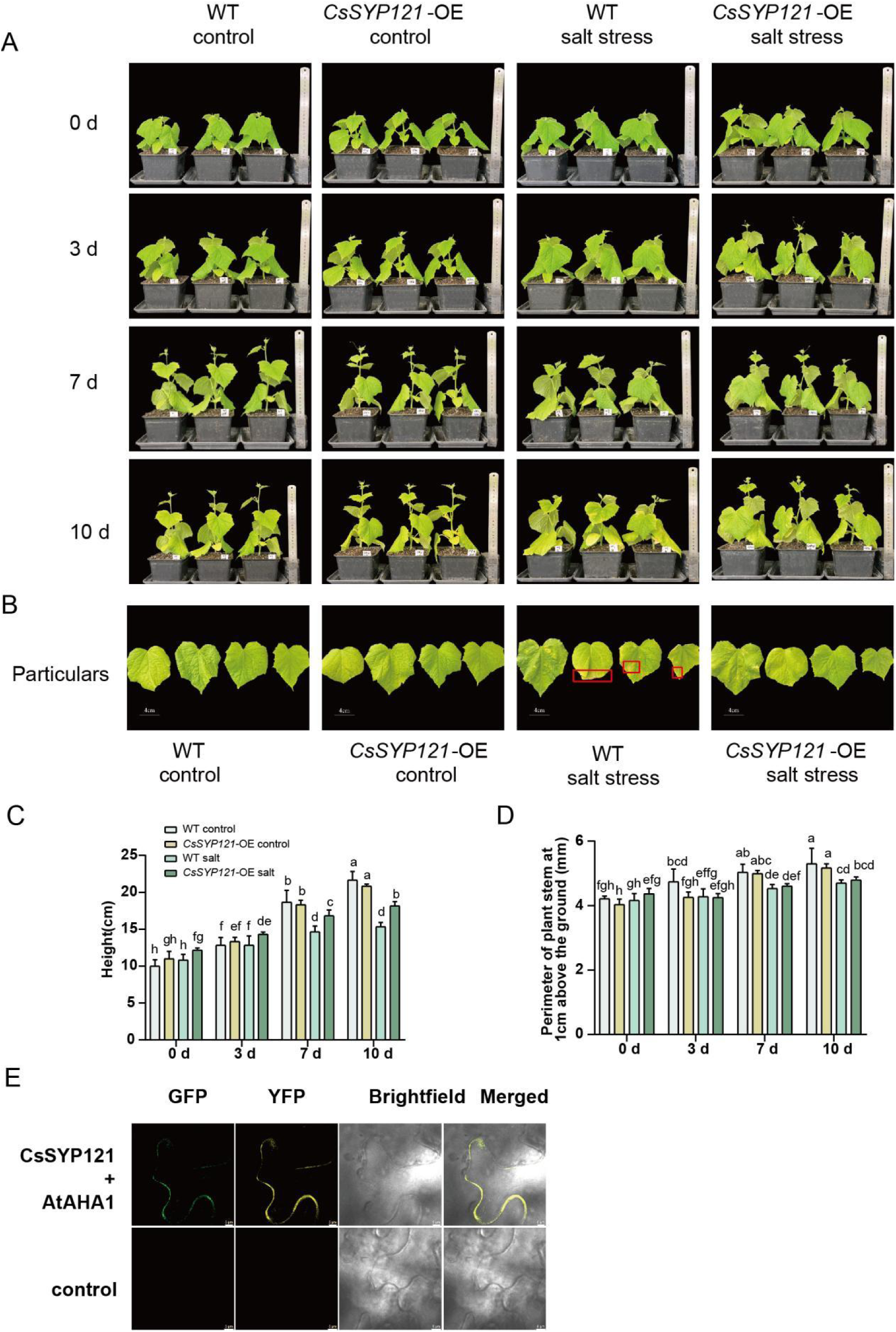
Comparison of salt tolerance between *CsSYP121*-OE and wild-types cucumber plants. (A) The growth phenotypes of *CsSYP121*-OE and wild-types cucumber plants under salt stress of 0, 3 and 7 days. (B) The growth phenotypes and details of leaves of *CsSYP121*-OE and wild-types cucumber plants under salt stress for 10 days. The red boxes indicate leaves with significant phenotypic differences. (C) The plant heights of *CsSYP121*-OE and wild-types cucumber plants under salt stress of 0, 3 and 7 days. (D) Stem thickness of *CsSYP121*-OE and wild-types cucumber plants under salt stress of 0, 3 and 7 days. (E) Subcellular localization of CsSYP121. Images from left to right are GFP fluorescence (CsSYP121), YFP fluorescence (AtAHA1), bright field, and merged field of view, respectively. Bright-field shows the tobacco cells. Merged image represents the overlay of the tw<u>o</u> channels. Scale bar = 5 μm. Results are from three independent experiments (biological replicates, n=3). Data points represent the mean of each independent experiment, with error bars showing SEM. Different letters indicate significant differences (*P*<0.05).

Next, we examined the effects of *CsSYP121* overexpression on water content and soluble solute content in cucumber plants under salt stress (Fig. 11). The results demonstrated that *CsSYP121* overexpression alleviated the increase in soluble solute content induced by salt stress. K^+^ and Na^+^, as critical inorganic salt ions, play vital roles in various key physiological processes, including osmoregulation in plants. In further experiments, we found that salt stress treatment resulted in increased Na^+^ content and decreased K^+^ content in the plants. However, *CsSYP121* overexpression effectively mitigated these alterations in ion levels. Within plant, ROS function as essential signaling molecules. However, their pathological accumulation frequently acts as a primary contributor to stress-elicited cellular damage. Consequently, strategies aimed at attenuating excessive ROS accumulation constitute a fundamental defense mechanism during stress responses (Ali et al., 2023; Foyer and Noctor, 2016). Enough K^+^ supply is important for reducing ROS-related damage induced by stress (Liu and Liao, 2022; Mostofa et al., 2022). Our prior work in foxtail millet demonstrated that high potassium levels alleviated leaf ROS accumulation triggered by nanoplastic treatment (Guo et al., 2024). Building upon these findings, we subsequently measured ROS levels and antioxidant enzyme activities in cucumber. Under control conditions, *CsSYP121* overexpression exhibited no significant effects H_2_O_2_ levels or antioxidant enzyme activities. Following salt stress imposition, wild-type plants experienced a burst of ROS generation leading to injury. In contrast, *CsSYP121* overexpressing plants displayed significantly enhanced antioxidant enzyme activity, which effectively counteracted the rise in ROS levels.

**Fig. 11.**
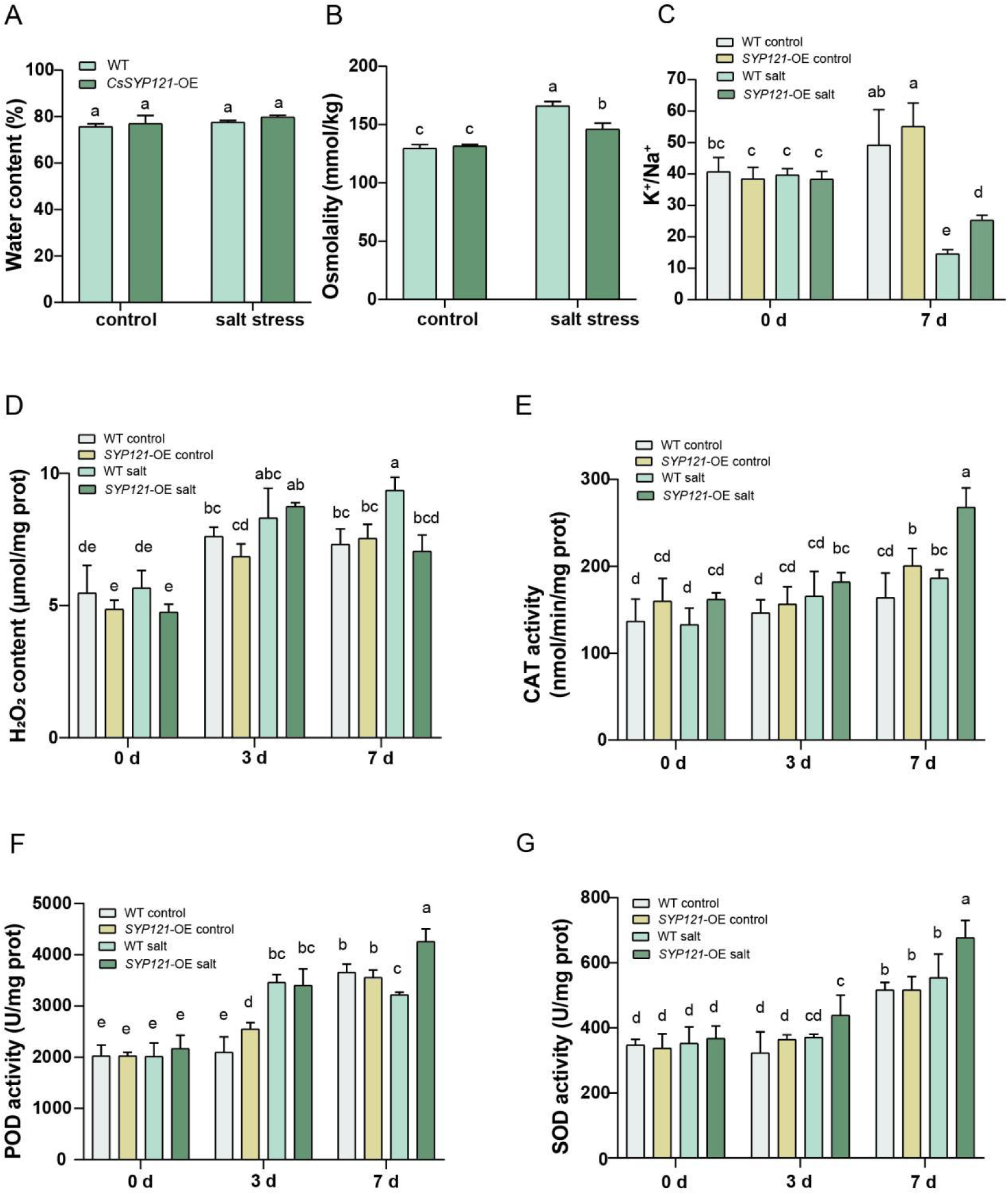
The effects of salt stress on water content (A), osmotic pressure(B), K^+^/Na^+^ content ratios (C), H_2_O_2_ activity (D), CAT activity (E), POD activity (F), and SOD activity (G) of SYP121-OE and wild-type cucumbers. Results are from three independent experiments (biological replicates, n=3). Data points represent the mean of each independent experiment, with error bars showing SEM. Different letters indicate significant differences (*P*<0.05).

## 4. Discussion

Throughout their life cycles, plants routinely encounter various biotic and abiotic stresses. Within plant systems, SNARE proteins represent a functionally diverse family exhibiting multifaceted biological roles, particularly in responding to both biotic and abiotic challenges (Gu et al., 2020; Zhang et al., 2020). Cucumber (*C. sativus* L.), as an agriculturally and nutritionally valuable species, possesses considerable significance in horticulture, economy, and human nutrition. In this study, we identified and characterized 51 members of the *CsSNARE* gene family within the cucumber genome. Comprehensive bioinformatic analyses were performed to examine their physicochemical characteristics, cis-acting elements, and expression patterns under abiotic stress conditions. Furthermore, we investigated the function of *CsSYP121*-a member of the *Qa-CsSNARE* subfamily-in salt stress responses through its overexpression in cucumber. Our findings demonstrate that *CsSYP121* overexpression mitigates excessive ROS accumulation by regulating K^+^ and Na^+^ levels, thereby contributing to cucumber’s salt stress adaptation.

Drawing upon genetic data from Arabidopsis and rice, this study identified 51 putative *CsSNARE* genes in cucumber—surpassing counts observed in algae (n=20), humans (n=35), and yeast (n=24)-while approximating the complement size in foxtail millet (n=52) yet remaining lower than that documented in Arabidopsis (n=65; Zhang et al. 2020). This expansion aligns with evidence suggesting increased *SNARE* gene dosage correlates with enhanced multicellular complexity during green plant evolution (Sanderfoot, 2007). Among these, we classified 14 *Qa-CsSNARE*, 9 *Qb-CsSNARE*, 10 *Qc-CsSNARE*, 3 *Qb+c-CsSNARE*, and 15 *R-CsSNARE* members (Fig. 1). Such conserved categorization patterns underscore the fundamental role of *SNARE* families in plant environmental adaptation. Comparative genomic analyses revealed significantly more orthologous gene pairs between cucumber and Arabidopsis than between cucumber and rice, attributable to their shared dicotyledonous lineage and correspondingly higher sequence conservation (Soltis et al., 2019).

Tissue-specific expression profiling demonstrated distinct spatial regulation within the *CsSNARE* family (Fig. 6). Notably, *Qa-CsSNARE* members including *CsSYP121*, *CsSYP41*, *CsSYP131b*, and *CsSYP122* exhibited root-enriched transcript abundance, implicating roles in root development—consistent with Arabidopsis *SYP121* function in root hair growth and functional redundancy between SYP121/SYP122 paralogs (Waghmare et al. 2018). Our observation of near-identical expression profiles for *CsSYP121* and *CsSYP122* across tissues supports this redundancy hypothesis. Conversely, most *Qb-CsSNARE* genes displayed elevated expression in meristematic and rapidly proliferating tissues (hypocotyls, seedling/adult stems, tendrils, shoot apex), mirroring the proliferation-associated expression pattern reported for *AtNSPN11* (Zheng et al., 2002) and suggesting potential functional overlap among family members. Distinctively, *Qc-CsSNARE CsSYP61* showed leaf-specific expression across developmental stages, coinciding with its proposed role in coordinating plant responses to carbon/oxygen dynamics (Hasegawa et al., 2022) and potentially regulating vegetative growth. Most members of the *R-CsSNARE* family exhibited higher expression levels in roots, stem, and shoot buds, while *CsVAMP722* and *CsSec223* were highly expressed across all tissues, including leaves. These collective findings validate our comprehensive identification of the *CsSNARE* gene family.

The cis-acting element analyses uncovered stress-responsive motifs (jasmonic acid responsive, low-temperature inducible, defense/stress-related) within *CsSNARE* promoters (Fig. 4). Prior studies have characterized specific SNARE complexes-such as the SYP121-SNAP33-VAMP721/722 complex-as components of Arabidopsis responses to biotic stresses (Kwon et al. 2008; Zhang et al. 2020). Analysis of publicly available RNA-Seq datasets revealed that most *CsSNARE* genes underwent rapid transcriptional activation within 24-48 hours following challenges by angular leaf spot pathogens and *C. cucumerium* infection. In contrast, Gray mold and Powdery mildew infections elicited elevated expression of these genes only after 3-5 days (Fig. 7), indicating functional diversification among *SNARE* family members during distinct biotic stress responses. Under abiotic stress conditions, heat and salt treatments broadly upregulated most *CsSNARE* genes, whereas cold stress induced significant expression changes in a subset comprising *CsSYP112*, *CsSYP122*, *CsSYP123*, *CsSNAP30*, *CsNPSN11*, *CsNPSN12*, and *CsVAMP721*.

SNARE proteins assume a central role in mediating plant responses to abiotic stressors-including drought, high salinity, and extreme temperatures (Pratelli et al., 2004). Through their function in membrane fusion, SNAREs facilitate the targeted transport of stress-responsive proteins, including antioxidant related proteins, to specific subcellular compartments, a process frequently accompanied by dynamic shifts in protein localization (Neves et al., 2021). In this study, promoter analysis of *CsSNARE* genes revealed that members of the *Qa-CsSNARE* subgroup harbor multiple cis-acting elements linked to drought responsiveness, including an MBS (MYB binding site) implicated in drought induction (Fig. 4). Consistent with this regulatory architecture, experimental evidence confirmed that *Qa-CsSNAREs* exhibit significant transcriptional activation under salt stress conditions (Fig. 5). Accordingly, our investigation centers on characterizing the stress-resistant functions of *Qa-CsSNAREs* in cucumber. Notably, *CsSYP41* displayed significant upregulation exclusively under drought stress, with minimal response to salt treatment (Fig. 8, 9). Supporting a functional role for SYP4 family members in stress adaptation, previous studies documented growth inhibition in *syp4* double mutants under osmotic and salt stress regimes (Uemura et al., 2012), proposing a mechanism involving protein complex formation. In contrast, *CsSYP121* expression remained largely unchanged under drought but showed marked upregulation under salt stress (Fig. 8, 9). Overexpression of *CsSYP121* did not confer discernible phenotypic alterations under drought conditions compared to wild-type plants (Fig. S1). Based on combined phenotypic and gene expression analyses under both stresses, we hypothesize that *CsSYP121* operates beyond general osmotic stress regulation, potentially modulating ionic homeostasis specifically under salt stress—a notion aligned with findings from previous reports(Eisenach et al. 2012; Besserer et al. 2012) . Further research is warranted to elucidate the signaling specificity underlying these distinct responses.

As a characterized ABA-responsive Qa-SNARE involved in stomatal regulation (Eisenach et al., 2012; Leyman et al., 1999), SYP121 emerged as a candidate regulator of salt stress responses in our study. We observed a significant induction of *CsSYP121* expression in cucumber leaves at 7 days post-salt treatment (Fig. 9). To test its functionality, we generated *CsSYP121*-overexpressing lines, which displayed enhanced salt tolerance compared to wild-type controls (Fig. 10). This result aligns with findings from soybean (Hong et al., 2022), where SYP121 was proposed to play a positive role in salt stress acclimation. These results reinforce a broader theme: manipulating SNARE-mediated membrane traffic appears to be a conserved strategy for improving salt tolerance across species, as evidenced by the effects of *GsSNAP33*, *AtSFT12*, and *SlSLSP6* overexpression (Nisa et al., 2017; Salinas-Cornejo et al., 2023; Tarte et al., 2015).

Our findings show that overexpression of *CsSYP121* has no significant impact on cucumber growth (Fig. 10). By contrast, Arabidopsis-based analyses indicate that *SYP121* overexpression confers a moderate enhancement to seedling development (Zhang et al., 2019), thereby demonstrating functional divergence of this gene across plant species. Our data suggest a complementary mechanism for *CsSYP121* function under salt stress. Previous research from Arabidopsis links SYP121 directly to ionic homeostasis. Recessive *syp121* mutants exhibit salt hypersensitivity due to impaired K^+^/Na^+^ balance (Eisenach et al., 2012). Molecularly, SYP121 partners with KC1 and AKT1 to promote K^+^ uptake (Honsbein et al., 2009) and also interacts with the KAT1 channel (Honsbein et al., 2011; Lefoulon et al., 2018). In our study, *CsSYP121*-OE plants maintained significantly higher K^+^ levels under salt stress (Fig. 11C) and reduced ROS accumulation (Fig. 11D-G). Maintaining a high cytoplasmic K^+^ concentration is crucial for counteracting Na^+^ toxicity, preserving enzymatic activity, and sustaining osmotic homeostasis (Fang et al., 2021). High intracellular K^+^ not only supports cell turgor but also helps counteract Na^+^ toxicity and reduces ROS accumulation (Che et al., 2024; Guo et al., 2024). Thus, *CsSYP121* likely enhances salt tolerance by improving K^+^/Na^+^ homeostasis and reducing ROS damage. Further studies are warranted to investigate potential interactions and regulatory relationships between *CsSYP121*, cucumber K^+^ channels, and Na^+^ transporters.

## 5. Conclusion

This study employed a genome-wide research to identify 51 *CsSNARE* genes in cucumber. Based on phylogenetic analysis, conserved motif characterization, and exon-intron structure, these genes were classified into five distinct subfamilies. Transcript profiling across diverse tissues, developmental stages, and stress treatments demonstrated that the *Qa-CsSNARE* subfamily is highly responsive to abiotic stress. Further examination pinpointed *CsSYP121* as a positive regulator of the salt stress response. To validate its function, we generated *CsSYP121*-overexpressing lines, which exhibited significantly enhanced salt tolerance compared to wild-type plants. Under salt stress, these transgenic plants showed improved ROS scavenging capacity and K^+^/Na^+^ homeostasis. In summary, this study not only systematically elucidates the evolutionary and expression characteristics of the *SNARE* gene family in cucumber but, more importantly, preliminarily reveals the molecular mechanism by which CsSYP121 enhances salt tolerance through the regulation of ion homeostasis and redox balance. This provides an important candidate gene and theoretical foundation for breeding salt-tolerant cucumber varieties using gene editing or molecular marker-assisted selection.

## CRediT authorship contribution statement

**Ben Zhang:** Writing - review & editing, Supervision, Software, Methodology, Funding acquisition, Data curation. **Weijuan Zhou:** Writing - review & editing, Writing - original draft, Validation, Supervision, Methodology. **Jie Zheng:** Validation, Software, Methodology, Investigation, Data curation. **Shengjun Zhou:** Validation, Investigation. **Yue Guo:** Validation, Investigation. **Dongmei Kong:** Visualization, Software, Methodology. **Pu Yang:** Investigation, Formal analysis.

## Declaration of Competing Interest

The authors declare that they have no known competing financial interests or personal relationships that could have appeared to influence the work reported in this paper.

## Acknowledgements

We thank all the colleagues in our laboratory for providing useful discussions and technical assistance. This work was supported by the National Natural Science Foundation of China (No. 32272012), the Fund Program for the Scientific Activities of Selected Returned Overseas Professionals in Shanxi Province (No.20230003) to BZ. The funding bodies played no role in the design of the study and collection, analysis, and interpretation of data and in writing the manuscript.

## Appendix A. Supporting information

Supplement table 1. Primers used in this study

Supplement table 2. Characteristics of the promoter region of *CsSNARE* genes in *Cucumis sativus* L. Supplement Fig. S1 The phenotypes of *CsSYP121*-OE and wild-type cucumber plants after 14 days of drought

Supplement Fig. S2 Under salt stress, the expression level of the *CsSYP121* gene in the roots of wild-type cucumbers.

Supplement Fig. S3 The expression level of *CsSYP121*-OE plants was validated by qRT-PCR.

Supplement Fig. S4 Western blot analysis using the CsSYP121 antibody was performed to verify the overexpression of CsSYP121 at the protein level.

## Data availability

The datasets generated during and analysed during the current study are available from the corresponding author on reasonable request.

## Notes

### Competing Interest Statement

The authors have declared no competing interest.

