## Supplement figure for "Identification of *SNARE* Genes in Cucumber and the Role of *CsSYP121* in Salt Stress Response": supplement figure20260227.docx

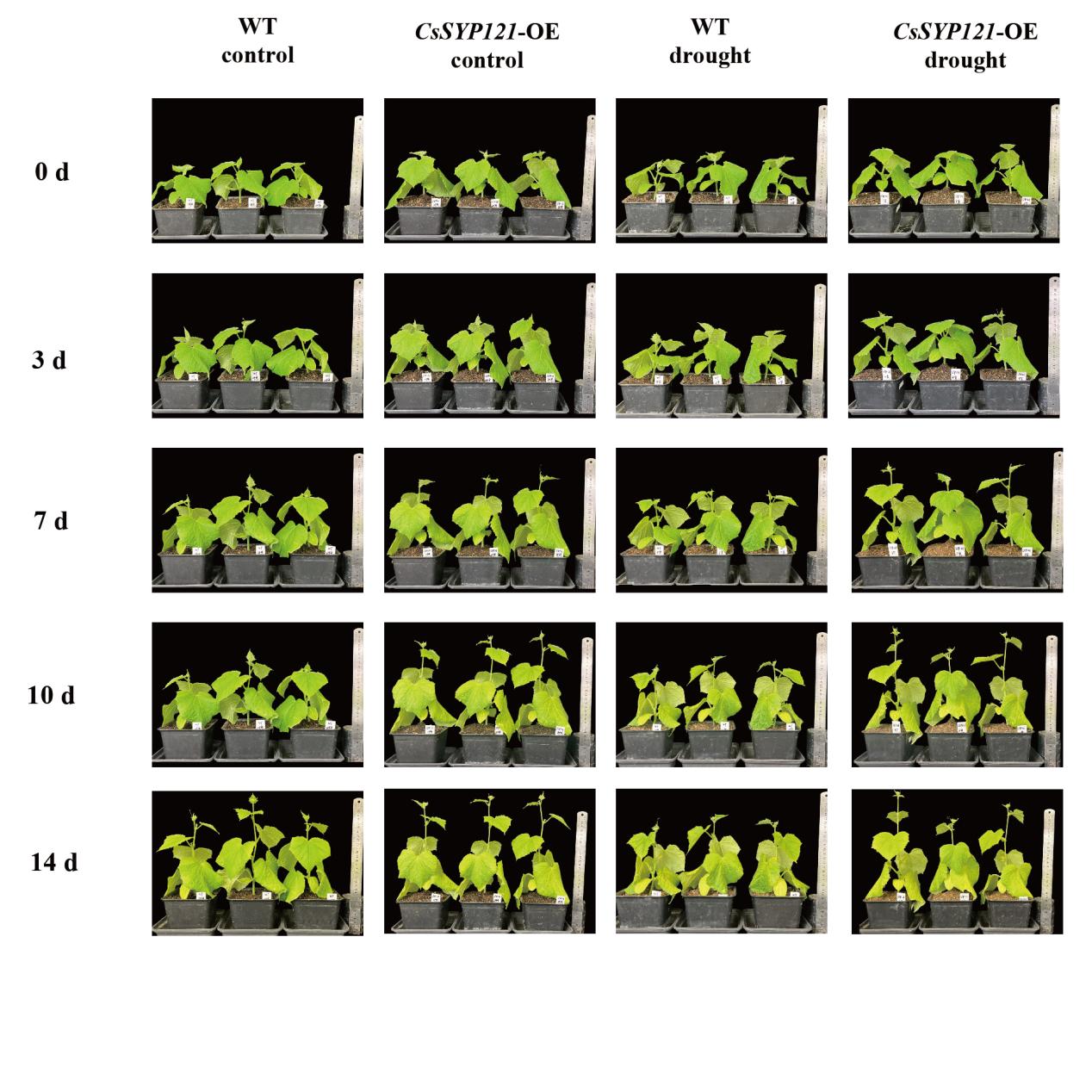

Fig. S1 The phenotypes of *CsSYP121*-OE and wild-type cucumber plants after 14 days of drought

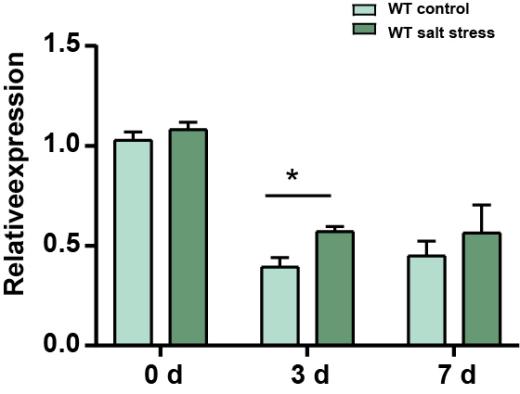

Fig. S2 Expression analysis of CsSYP121 in wild-type cucumber roots under salt stress. Transcript levels of CsSYP121 were examined by qRT-PCR in roots of wild-type plants under control conditions or after 3 and 7 days of salt treatment. Each bar represents the mean ± SE normalized to *CsEF1α* (*CsaV4_5G001996*) and *CsCACS* (*CsaV4_3G004932*). Three biological and three technical replicates were performed for each sample. Light green and dark green columns indicate wild-type plants under control and salt stress conditions, respectively. Asterisks indicate statistically significant differences compared with the control (**P* < 0.05, ***P* < 0.01, ****P* < 0.001).

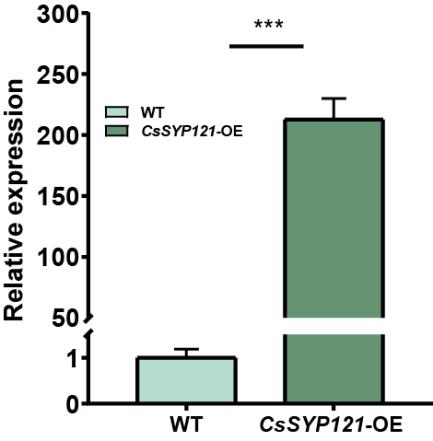

Fig. S3 The expression level of *CsSYP121*-OE plants was validated by qRT-PCR. When the cucumber seedlings reached three leaves and one heart stage (three fully expanded cotyledons and visible apical meristem), the plants with consistent growth condition were selected for sampling. Each bar represents the mean±SE normalized to *CsEF1α* (*CsaV4_5G001996*) and *CsCACS* (*CsaV4_3G004932*). All samples were run in three biological and three technical replicates. Light green columns represent “WT”, and Dark green columns represent “*CsSYP121*-OE”. Asterisk indicates that the gene expression after stress has a significant difference compared with the control (****P*<0.001).

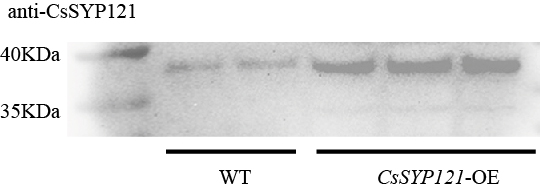

Fig. S4 Western blot analysis using the CsSYP121 antibody was performed to verify the overexpression of CsSYP121 at the protein level.
